# Human monoclonal antibodies that target clade 2.3.4.4b H5N1 hemagglutinin

**DOI:** 10.1101/2025.02.21.639446

**Authors:** Garazi Peña Alzua, André Nicolás León, Temima Yellin, Disha Bhavsar, Madhumathi Loganathan, Kaitlyn Bushfield, Philip J.M. Brouwer, Alesandra J. Rodriguez, Trushar Jeevan, Richard Webby, Christine Marizzi, Julianna Han, Andrew B. Ward, J. Andrew Duty, Florian Krammer

## Abstract

The highly pathogenic avian influenza H5N1 virus clade 2.3.4.4b has been spreading globally since 2022, causing mortality and morbidity in domestic and wild birds and mammals, including infection in humans, raising concerns about its pandemic potential. We aimed to generate a panel of anti-hemagglutinin (HA) human monoclonal antibodies (mAbs) against the H5 protein of clade 2.3.4.4b. H2L2 Harbour Mice^®^, which express human immunoglobulin germline genes, were immunized with H5 and N1 recombinant proteins from A/mallard/New York/22-008760-007-original/2022 H5N1 virus, enabling the generation of human chimeric antibodies. Through hybridoma technology, sixteen full human mAbs were generated, most of which showed cross-reactivity against H5 proteins from different virus variants. The functionality of the sixteen mAbs was assessed *in vitro* using hemagglutination inhibition and microneutralization assays with viruses containing a clade 2.3.4.4b HA. Fourteen out of the sixteen mAbs neutralized the virus *in vitro*. The mAbs with the strongest hemagglutination inhibition activity also demonstrated greater neutralizing capacity and showed increased protective effects *in vivo* when administered prophylactically or therapeutically in a murine H5N1 challenge model. Using cryo-electron microscopy, we identified a cross-clonotype conserved motif that bound a hydrophobic groove on the head domain of H5 HA. Akin to mAbs against severe acute respiratory syndrome coronavirus 2 (SARS-CoV-2) during the coronavirus 2019 (COVID-19) pandemic, these mAbs could serve as important treatments in case of a widespread H5N1 epidemic or pandemic.

## Introduction

Highly pathogenic H5N1 avian influenza virus emerged as a human disease in 1997^1^. After variable H5N1 virus activity in the years before the coronavirus disease 2019 (COVID-19) pandemic, a subclade of H5N1, clade 2.3.4.4b, started to spread globally in 2022. This spread has resulted in significant losses in the poultry industry and endangered millions of wild birds^2,3^. This virus clade has also spilled over into mammals, causing severe disease and high fatality rates^4^. In March 2024, the virus was detected in the United States (U.S.) dairy cattle, spreading among herds with high titers of virus present in the cow’s milk and in October 2024, virus was detected in a pig in a backyard farm in Oregon^5^. Even more alarming is the reported transmission of the virus to humans, posing a risk to workers who are in contact with infected animals and raising concerns about the potential for a pandemic. So far, more than 70 clade 2.3.4.4b human infections have been reported, with the vast majority in the U.S. While antivirals licensed for seasonal influenza are likely effective for H5N1, there are currently no long-acting treatments available^6,7^. To address this issue, we have developed human monoclonal antibodies (mAbs) generated in immunoglobulin germline humanized mice. These mAbs can effectively neutralize clade 2.3.4.4b H5N1 and provide protection in a murine H5N1 challenge model. MAbs have been used extensively and successfully as therapeutics and prophylactics during COVID-19 pandemic^8,9^ the mAbs developed here against H5N1 may play roles as therapeutics or prophylactics in potential future H5N1 epidemics or pandemics.

## Results

### Generation and *in vitro* characterization of mAbs

H2L2 Harbour Mice^®^, which express human-rat chimeric antibody genes^10^, were immunized with recombinant hemagglutinin H5 and neuraminidase N1 proteins of clade 2.3.4.4b virus (Supplementary Fig.1A) and mAbs were generated via hybridoma technology. Initial screening was done using enzyme-linked immunosorbent assays (ELISA) and hemagglutination inhibition (HI) assays, identifying 1,977 positive clones. From these, 401 clones were down-selected for secondary screening, of which 30 clones showed strong binding to the H5 protein and had HI activity, leading to their selection for isotyping and sequencing. After removal of IgMs and identical clones, down-selected clones were chosen for variable segment DNA synthesis and cloning into human constant containing-expression vectors (human G1/ kappa) for fully human recombinant antibody production. Sixteen recombinant clones were selected as sequence unique lead candidates based on broad H5 activity and recombinant expression. We grouped these sixteen antibodies into clonotypes based on exact amino acid length with >90% amino acid identity in their junctions/complementarity-determining 3 (CDR3) regions along with shared V(D)J/VJ gene usage for both their heavy and light chains, resulting in a total of 5 closely related clonotypes (Supplementary Fig.1D). The identified clonotypes and their corresponding clones were as follows: clonotype 1 (1A1, 12C11, 12G9, 13B9, 13E8), clonotype 2 (1H2, 17E3, 20D10), clonotype 3 (6G1, 7G4, 7G11, 12G8, 17C12), clonotype 4 (7H10, 14B8) and clonotype 5 (12G1). While clonotypes 3, 4 and 5 each utilized the longest heavy chain CDR3s at 19 amino acids (IMGT numbering) and had shared heavy V(D)J usage (VH4-59, DH3-10, and JH6), the sequence variation in their CDR3 was greater than 40% which was significantly higher than the predicted somatic hypermutation rates (1.7-3.6%) when analyzing their VH4-59 variable genes alone to germline (data not shown). This argues for separate clonal ancestries and, thus, their categorization as unique clonotypes. Light chain CDR3 lengths were less varied given the lack of a D gene cluster and averaged 9 amino acids for each clonotype, typical for human kappa chains, even though clonotype 4’s light chain was one residue shorter at 8 amino acids, another unique characterization for this clonotype.

These mAbs were expressed, and their binding activity was assessed against a panel of H5 proteins from virus of different clades (Fig.1A). The mAbs showed robust binding to H5 HAs from recent clade 2.3.4.4b isolates including to HAs from viruses recently detected in birds in New York City, New York, U.S.^11^. Of note, robust binding to the HA of a U.S. bovine H5N1 isolate was detected as well. Most mAbs also showed strong binding to HAs from older clade 2.3.4.4 isolates. This included avian strains from 2014 from the Netherlands and the U.S. and an H5N6 human isolate from a 2016 case in China. However, no binding was observed for HAs from older clade 1 from 2004 and 2 strains, for HA from the 2.3.2.1c clade that has been causing recent human cases in Cambodia or for H1 HA from seasonal H1N1 strain (used as negative control). The phylogenetic relationship of these HA proteins is shown in Supplementary Fig.2. Most clonotypes, except for the one containing mAbs 7H10 and 14B8, showed strong binding to H5 of clade 2.3.4.4b H5N1 viruses. Clones belonging to the same clonotype typically exhibited similar binding patterns. Further characterization of the mAbs was performed *in vitro* using HI and microneutralization assays. An HI assay was run to assess whether the mAbs inhibit the interaction of the virus with the host receptor (Fig.1B). This assay was conducted with several virus strains. Antibodies showed HI activity against A/bald eagle/FL/W22-134-OP/2022 (H5N1-A/PR/8/34 reassortant), here referred to as A/bald eagle/FL/W22-134-OP/2022 and viruses isolated from dairy cattle (clade 2.3.4.4b also) in a similar manner. As expected, in accordance with the binding data, no HI activity was observed against the A/Vietnam/1204/2004 virus belonging to clade 1. A/Victoria/4897/2022, a human seasonal H1N1 was used as irrelevant virus control. No HI activity was detected against this strain demonstrating specificity to clade 2.3.4.4 H5N1 viruses. Of note, the clonotype containing clones 7H10 and 14B8, not only had low binding, but also showed no HI activity. Additionally, an *in vitro* microneutralization assay was performed to assess the ability of the mAbs to disrupt the viral lifecycle, potentially through various mechanisms (Fig.1C). We tested whether the antibodies showed potential to prevent the virus from entering the cell or to block the virus release, or both. The same fourteen mAbs with the highest HI activity also exhibited greater neutralizing capacity and inhibited viral replication in cell culture at different stages of the virus life cycle. The mAbs 1A1 from clonotype 1, as well as all the clones belonging to clonotype 2, 3, and 5 with the strongest HI activity, demonstrated greater neutralizing capacity, with a minimal neutralizing titer of up to 0.024ng/ml being observed. As the final assay, we assessed whether the antibodies inhibited the neuraminidase (NA) enzymatic activity through steric hindrance. Clone 1H2 from clonotype 2 and 12G1 from clonotype 5 showed strong inhibitory activity, similar to the previously characterized pan-NA antibody, 1G01^12^, here used as a positive control. 7H10 and 14B8 from clonotype 4 exhibited low or non-NA inhibitory activity.

**Figure 1.**
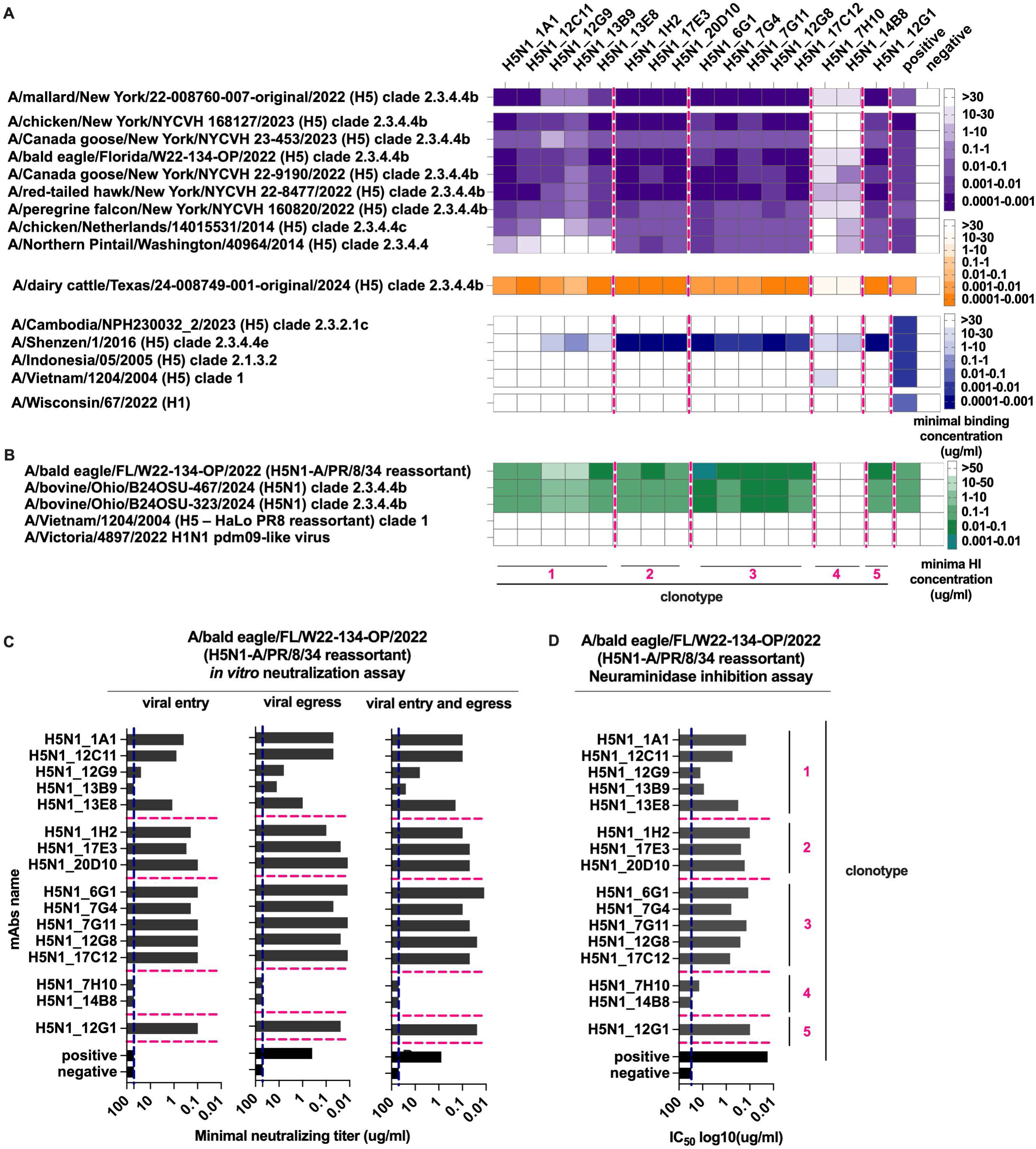
Anti-H5 mAbs bind broadly to HA proteins of virus variants and inhibit virus replication *in vitro*. **(A)** Binding breath of anti-H5 mAbs to H5 protein from infected birds (purple), cattle (orange), and individuals (blue), was evaluated by ELISA in duplicates. Positive control an anti-HA antibody, CR9114^27^. Data is minimal binding concentration (ug/ml), defined as the concentration with a signal greater than 3 standard deviations above blanks. **(B)** mAbs ability to block the virus-host receptor interaction was assessed by hemagglutination inhibition (HI) assay in duplicates. Positive control, serum (1:10) from a mouse infected with A/bald eagle/FL/W22-134-OP/2022 (H5N1-A/PR/8/34 reassortant). Data shows minimal HI concentration (ug/ml), defined as the last concentration that neutralizes the virus. **(C)** Efficacy of the anti-H5 mAbs to inhibit viral replication was evaluated using an *in vitro* neutralization assay against A/bald eagle/FL/W22-134-OP/2022 (H5N1-A/PR/8/34 reassortant), in duplicates, with controls identical to part A. Data shows minimal neutralizing titer (ug/ml), defined as the last concentration where neutralization is observed. **(D)** mAbs capacity to inhibit neuraminidase (NA) activity of clade 2.3.4.4b virus was tested in duplicates, with positive control an anti-NA antibody, 1G01^12^. Data shown half inhibitory concentration (IC_50_) (ug/ml). Negative control for all the assays was an anti-SARS-CoV-2 spike antibody^13^.

### Human anti-H5 mAbs provide protection in a murine H5N1 challenge model when given prophylactically or therapeutically

To further investigate the antiviral activity of the mAbs *in vivo*, we tested select mAbs from different clonotypes in both prophylactic and therapeutic settings in an H5N1 challenge model in 6-week-old BALB/c mice. The mAbs with the strongest binding and highest neutralization activity from each clonotype, 1A1 (clonotype 1), 20D10 (clonotype 2), 6G1 (clonotype 3) and 12G1 (clonotype 5), were selected for the *in vivo* studies. Mab 7H10 (clonotype 4) was added as representative for a binding but non-neutralizing mAb. In the prophylactic study, the mAbs were administered at four different doses via the intraperitoneal route, 4 hours prior to intranasal challenge with 50% lethal doses (LD_50_) of A/bald eagle/FL/W22-134-OP/2022. Mice were monitored for weight loss and survival for 14 days to assess protection (Fig.2A-C). The antibodies that were effective neutralizers *in vitro*, 1A1, 20D10, 6G1 and 12G1, also demonstrated the ability to protect mice in a dose-dependent manner. Mice administered with 5mg/kg, 1mg/kg, and 0.3mg/kg of the indicated antibodies showed no morbidity and complete survival. Of note, mice injected with antibodies belonging to clonotype 1 (1A1) and clonotype 3 (6G1) even demonstrated protection with a very low dose of 0.1mg/kg. This level of protection was not observed in mice administered with the non-neutralizing mAb 7H10 or with an isotype control antibody (hG1/hk, anti-severe acute respiratory syndrome coronavirus 2 (SARS-CoV-2) spike mAb^13^), indicating that protection in this case is associated with the administration of neutralizing antibodies.

**Figure 2.**
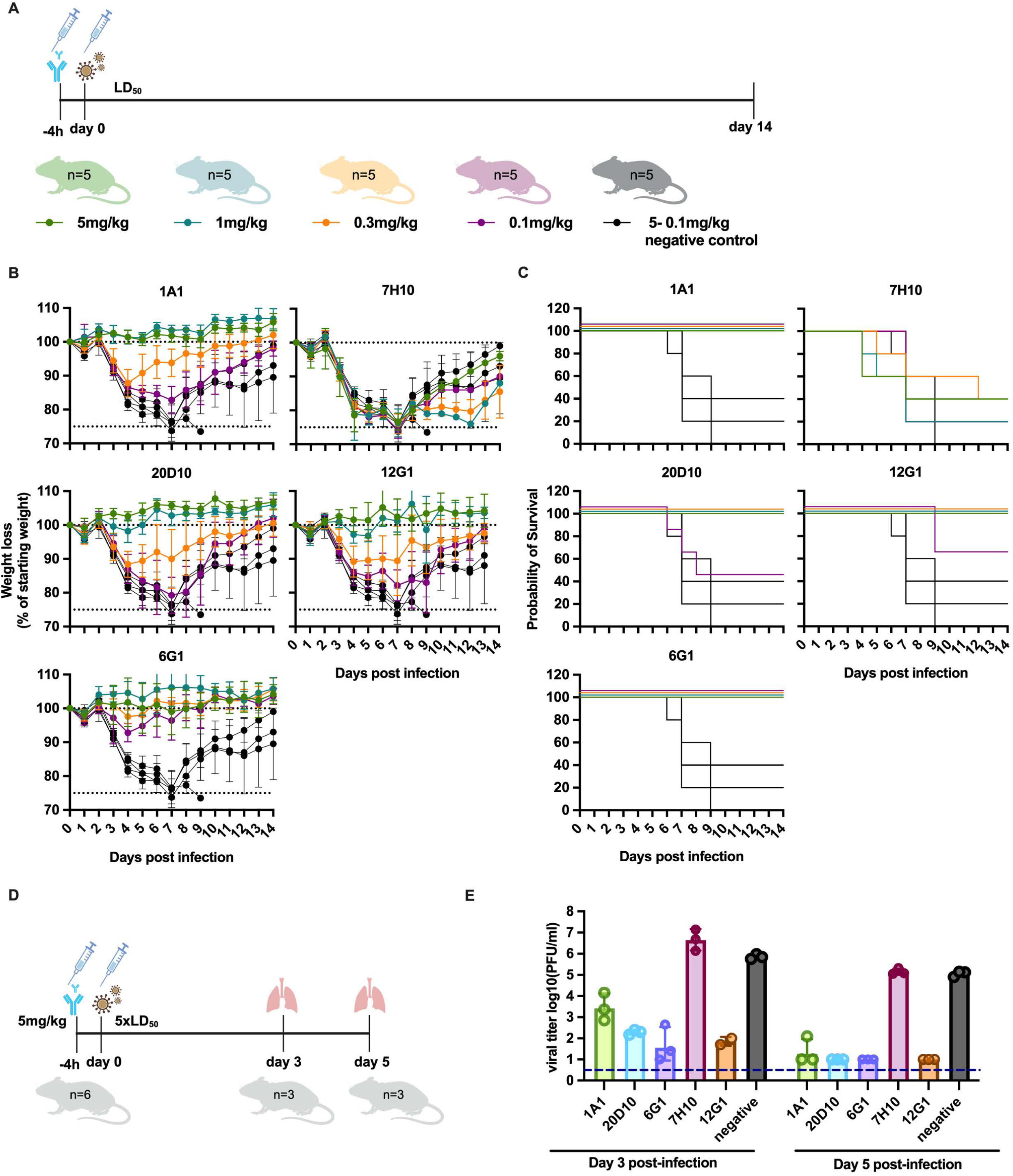
Anti-H5 mAbs are protective in a prophylactic setting *in vivo* and promote viral clearance in infected mice lungs. **(A)** Six-week-old female BALB/c mice (n=5/group) were injected intraperitoneally with different amounts of anti-H5 mAb, as a negative control anti-SARS-CoV-2 spike mAb was used, the data of negative control is plotted alongside the experimental groups. After 4h, mice were infected with 50% lethal doses (LD_50_) of A/bald eagle/FL/W22-134-OP/2022 (H5N1-A/PR/8/34 reassortant) virus. Body weight percentage **(B)** and survival **(C)** were plotted over 14 days post-infection. The percentage of the body weight is relative to the starting body weight, data is represented as group means with standard deviation. **(D)** Six-week-old female BALB/c mice (n=3/group) received 5mg/kg of anti-H5 mAbs, and anti-SARS-CoV-2 mAb was used as a control, 4h before infection done as in Part A. Lungs were harvested on day 3 and day 5 post-infection. **(E)** Viral titers in lungs from individual mice were determined as log_10_ plaque forming units (PFU)/ml/. Dots represent values from individual mice, and bars represent geometric means with geometric standard deviations. The dashed line at the Y-axis represents the limit of detection (LOD), defined by half of the lowest dilution at which a positive assay response is observed.

We then wanted to study if these mAbs could reduce viral replication in the lungs of animals. Experiments were performed similarly to the one described above with mAbs dosed 4 hours prior to challenge with 5 LD_50_ of A/bald eagle/FL/W22-134-OP/2022 virus. Lungs were harvested on days 3 and 5 post-infection, and viral titers were determined using plaque assay (Fig.2D, E). Viral titers were observed to be 4 logs lower in lung homogenates of mice treated with mAbs 20D10, 6G1 or 12G1 3 days post-infection, compared to the isotype control group. At day 3, we also observed a significant (3 log) reduction in viral titers in the lung homogenates of mice that received the 1A1 antibody. At day 5, no virus was detectible in the lungs of 20D10, 6G1 and 12G1 treated animals. No reduction was observed for mAb 7H10 compared to the control on day 5. This finding suggests that the antibodies inhibit replication of H5N1 in the lungs, which could be associated with their high neutralizing efficiency in cell culture.

To further assess the potential therapeutic benefits of the mAbs, mice were intranasally challenged with 5 LD_50_ of A/bald eagle/FL/W22-134-OP/2022 virus. MAbs were subsequently administered on days 2, 3, or 4 post-infection, and the mice were monitored for weight loss and survival for 14 days to evaluate the protective effect (Fig.3). All mice that received 6G1 mAb on day 2 post-infection survived, and 80% of mice that received 20D10 and 12G1 at the same time point survived as well. Only 40% of mice that received mAb 1A1 at day 2 post-infection survived. Mice that received 20D10, 6G1, and 12G1 antibodies on day 3 and day 4 post-infection, when significant weight loss had already occurred, showed varying levels of survival, ranging from 20% to 60%. All the mice treated with 7H10 and anti-SARS-CoV-2 reached the ethical endpoint on days 5 and day 6 post-infection, consistent with data from the prophylactic treatment study.

**Figure 3.**
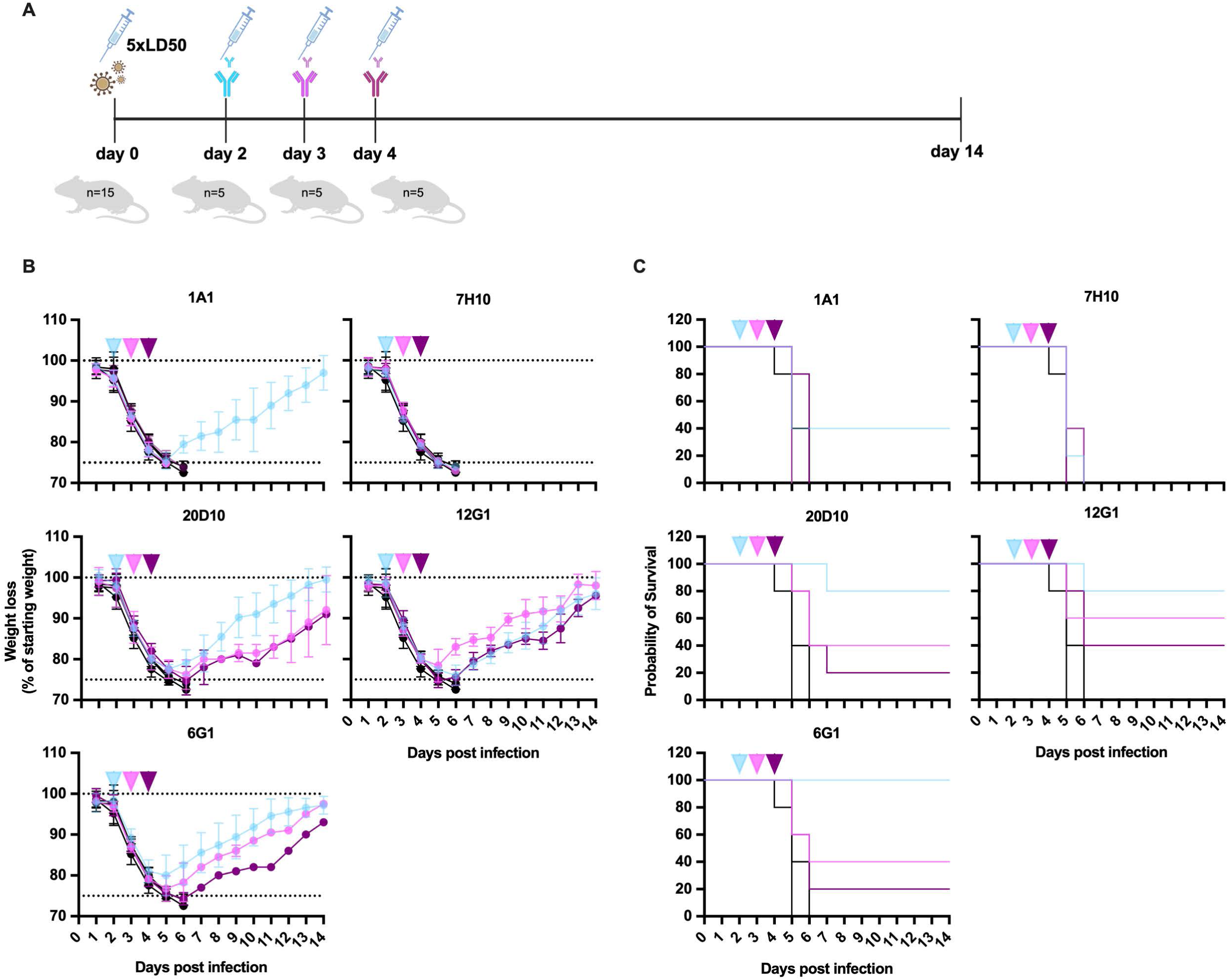
Anti-H5 mAbs are protective in a therapeutic setting *in vivo.* **(A)** Six-week-old female BALB/c mice (n=5/group) were injected intraperitoneally with 5mg/kg of anti-H5 mAb and as a negative control anti-SARS-CoV-2 spike mAb was used, the data of negative control is plotted alongside the experimental groups. The antibodies were administered on day 2, 3, and 4 after infecting the mice with 5 LD_50_ of A/bald eagle/FL/W22-134-OP/2022 (H5N1-A/PR/8/34 reassortant) virus. The percentage of initial body weight **(B)** and survival **(C)** were plotted over 14 days post-infection. The percentage of the body weight is relative to the starting weight, and the data is represented as group means with standard deviation. Orange arrows indicate the time point at which mAbs were administered.

### HA head regions are targeted by antibody clonotypes

Negative stain electron microscopy revealed that 13 representative mAbs from clonotypes 1-3 and 5 had binding footprints overlapping (clonotype 1) or proximal to the receptor binding site (RBS) (clonotypes 2, 3, and 5) when complexed with recombinantly expressed HA from A/Jiangsu/NJ210/2023 H5N1 (Jiangsu H5), a clade 2.3.4.4b clinical isolate from an infected human patient in Jiangsu, China (Supplementary Fig.3-4). Representative mAbs from clonotypes 1-3 and 5 were also complexed with recombinant HA from A/red-tailed hawk/New York/NYCVH 22-8477/2022, another clade 2.3.4.4b virus HA, indicating that the binding footprint remained consistent across strains (Supplementary Figs.3, 5). To assess the molecular basis of antigen engagement, we solved three high resolution cryo-EM structures from complexes with mAbs 20D10 (clonotype 2) at 3.4 Å resolution, 6G1 (clonotype 3) at 3.2 Å resolution, and 12G1 (clonotype 5) at 3.06 Å resolution (Supplementary Fig 6, Supplementary Table 1). Also, the 6G1 cryoEM sample was co-complexed with mAb 1A1 (clonotype 1); while the 1A1 density was insufficient for model building, we were able to dock a predicted model of the mAb to mark an approximate footprint. Our data revealed that clonotypes 2, 3, and 5 had nearly identical binding footprints proximal to the protomer interface while clonotype 1 had a distinct footprint that encompasses the RBS (Fig.4A). Visual inspection of the overall architecture of 20D10, 6G1, and 12G1 complexes revealed that epitope, angle of approach, and binding mode were structurally conserved across all three models (Fig,4B,C). Epitope contact analysis of interacting residues revealed that all three mAbs utilize a conserved LxYxY motif in CDRH3 to insert into a hydrophobic groove composed of residues 123-129 and 165-171 on Jiangsu H5, particularly an isoleucine-rich pocket composed of residues I123, I124, I164, and I166, which run antiparallel to one another (Fig.4D,E, Supplementary Table 2). In 20D10 and 6G1, these interactions are further stabilized by van der Waals interactions between D101 (mAb) and N129 (HA) on Jiangsu H5. Through sequence conservation analyses, we found that the hydrophobic groove is well conserved across the clade 2.3.4.4 strains in this study, but not amongst A/Wisconsin/67/2022 H1 or H5 clades 1, 2.3.2.1c and 2.1.3.2 (Fig.4F); amongst the identified interacting residues, I166 is particularly variable. While engagement with the hydrophobic groove was almost exclusively mediated by residues in the CDRH3, van der Waals and hydrogen bonding interactions were also identified between Jiangsu H5 and the CDRL1s and CDRL2s of all three mAbs (Fig.4G).

**Figure 4.**
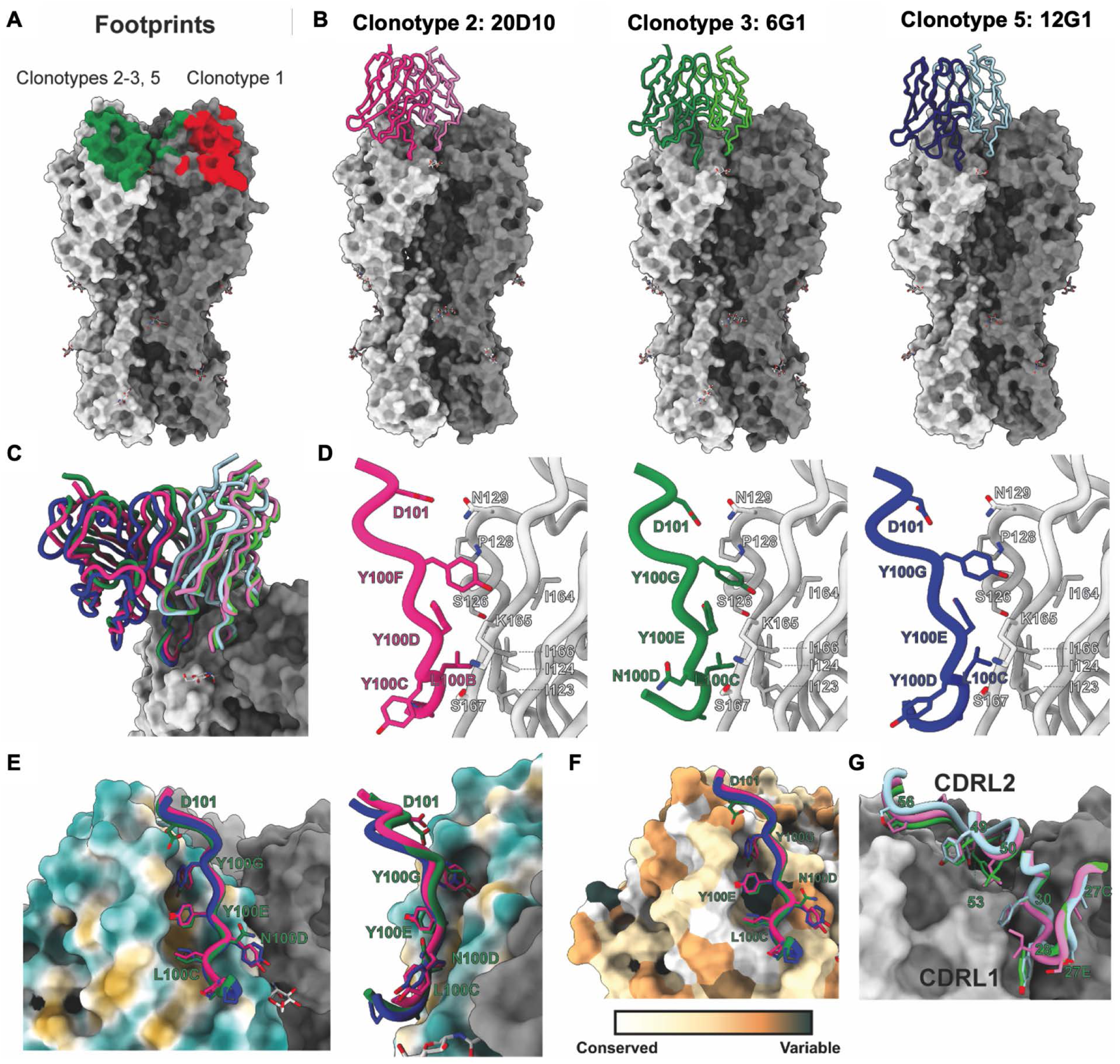
Structural characterization reveals shared approach to antigen engagement. **(A)** Footprints for clonotype 1 (red) and clonotypes 2, 3, and 5 (green) on a surface rendering of A/Jiangsu/NJ210/2023 H5. **(B)** Overall architecture of antigen engagement for mAbs 20D10 (pink), 6G1 (green), and 12G1 (blue) colored by clonotype. Heavy chains are presented in a darker shade than light chains. **(C)** Zoomed in overlay of heavy and light chain models of 20D10, 6G1, and 12G1 engaging with the antigen binding pocket. **(D)** Analysis of residues in mAb CDRH3 loops and the antigen binding pocket highlights conserved motifs across distinct clonotypes. Residue labels are presented in Kabat numbering and are colored by clonotype. **(E)** Hydrophobic surface renderings of the binding pocket and CDRH3 residues involved in direct engagement. Residue labels are provided using 6G1 numbering. **(F)** Amino acid sequence conservation of strains presented in Figure 1A mapped onto a surface rendering of A/Jiangsu/NJ210/2023 H5. Residue variability increases from white to orange to dark blue. **(G)** Antigen interactions mediated by CDRL1 and CDRL2 for mAbs 20D10, 6G1, and 12G1.

## Discussion

Here, we developed a panel of sixteen human antibodies against clade 2.3.4.4b highly pathogenic avian influenza (HPAI) H5N1 virus using an antibody germline humanized mouse system. Most of the generated antibodies showed broad binding within clade 2.3.4.4b HAs and many even bound to HAs from historic clade 2.3.4.4 strains. However, no cross-binding to older H5 clades or H1 HA was detected. Most mAbs showed strong *in vitro* antiviral activity in HI, neutralization and neuraminidase inhibition assays. Importantly, the HI active antibodies also had strong HI activity against two recent dairy cattle isolates. *In vivo*, all tested mAbs with *in vitro* neutralizing activity showed prophylactic activity against a lethal H5N1 challenge and also provided a survival benefit when given as late as 4 days post challenge, except for mAb 1A1 (clonotype 1) which only provided protection prophylactically. Structural characterization revealed that mAbs from clonotypes 2, 3 and 5 used a conserved motif to bind a hydrophobic groove on the HA head adjacent to the promoter interface while the clonotype 1 mAb bound directly to the RBS. So far in the U.S. more than 70 humans have been infected with clade 2.3.4.4b H5N1 virus with 6 severe infections^14–16^ including one mortality ^17,18^. While no human-to-human spread has been detected so far, mammalian adaptations have been found in clade 2.3.4.4b viruses that could increase the pandemic potential of the virus. In addition, a reassortment event between seasonal H1N1 or H3N2 with H5N1 could lead to the emergence of a virus that is pathogenic and transmissible in humans (of note, the last three influenza pandemics were caused by such reassortants)^19^. The antibodies described here, especially mAbs 12D10, 6G1 and 12G1, could be used as therapeutic treatment for severe zoonotic human H5N1 cases on their own or in addition to other antivirals like neuraminidase or endonuclease inhibitors^20,21^. Since 12D10, 6G1 and 12G1 target the same footprint, it is unlikely that using them together in a cocktail would be beneficial. But, especially for prophylactic treatment, one of them could be combined with mAb 1A1 which targets a different footprint and has shown potent protection when given prophylactically. These mAbs could also be combined with anti-influenza mAbs with other targets e.g. in the stalk domain of HA^22^ or on the viral NA^23^. In case of an H5N1 pandemic, it is likely that vaccines would be rolled out fast and that short acting antivirals like neuraminidase or cap-snatching inhibitors would be available.

However, similar to COVID-19, it is likely that a proportion of the population with underlying diseases or those that are immunocompromised will not be able to mount a sufficient response to vaccination and these individuals can also not be kept on antivirals for long periods of time. A long-acting mAb prophylaxis, as used for COVID-19 or RSV based on mAbs like 12D20, 6G1 or 12G1 could be used to protect vulnerable individuals during an H5N1 pandemic.

## Material and Methods

Most sections are further detailed in online supplemental materials.

### Cells

Madin-Darby canine kidney (MDCK) cells (ATCC #CCL-34) were used for *in vitro* microneutralization assays and viral titer determination. Expi293F cells were used for antibody production.

### Recombinant glycoproteins

Recombinant hemagglutinin (HA) and neuraminidase (NA) proteins were produced using the baculovirus expression system as described in^24^. References of the proteins from the indicated virus isolates, such as accession GISAID numbers and GenBank numbers, are indicated in the supplementary material. HA-expressing baculoviruses were passaged in Sf9 cells and then used to infect BTI-TN-5B1-4 cells for protein production. Proteins were purified using Ni2+-nitrilotriacetic acid (NTA) agarose beads (QIAGEN) three days post-infection, as described in^24^.

### Viruses

A/bald eagle/FL/W22-134-OP/2022 is a 6:2 reassortant H5N1 virus with A/PR/8/1934 rescued via reverse genetics, with a monobasic cleavage site replacing the polybasic cleavage site^25^. Similarly, A/Vietnam/1203/2004 virus retains surface H5N1 glycoproteins from A/Vietnam/1203/2004 (with a monobasic cleavage site replacing the polybasic cleavage site in the HA) while using the A/PR/8/34 backbone. The A/bovine/Ohio/B24OSU-467/2024 and A/bovine/Ohio/B24OSU-323/2024 are authentic bovine viruses.

### Immunizations and hybridoma generation

To generate human antibodies, H2L2 Harbour Mice^®^, human antibody transgenic mice (Harbour BioMed, Cambridge, MA) were used in collaboration between the Icahn School of Medicine at Mount Sinai and Harbour BioMed. The H2L2^®^ is a chimeric transgenic mouse containing the a selected human variable gene segment loci of the heavy and kappa antibody chains along with the rat heavy and kappa constant gene segment loci, producing antibodies with similar diversity seen in human antibody immunity^10,26^. Eight-to twelve-week-old H2L2 Harbour Mice^®^, mice immunized interperitoneally with HA and NA from A/mallard/New York/22-008760-007-original/2022 (H5N1) adjuvanted with 40ug of poly I:C (InVivoGen, San Diego, CA). Each mouse received a prime followed by two boosts, and blood was collected two weeks after each boost to monitor the titer of serum antibodies via enzyme-linked immunosorbent assay (ELISA) and hemagglutination inhibition (HI) analysis. One mouse was selected for hybridoma fusion and received two final boosts consisting of 20ug of the H5 protein at -4 and -2 days before being bled, euthanized by Institutional Animal Care and Use Committee (IACUC) approved methods, and spleen harvested for splenic fusions. The spleen was processed to a single-cell suspension, and hybridomas were generated using a standard protocol (StemCell Technologies, Vancouver, BC, Canada). Briefly, individual B cell clones were grown on soft agar and selected for screening using a robotic ClonaCell Easy Pick instrument (Hamilton/Stem Cell Technology). Individual clones were expanded, and the supernatant was used to screen for H5 binding (ELISA) and HI activity. All animal studies were approved by the Icahn School of Medicine (IACUC).

### Sequencing and humanizing of antibodies

Sequencing was done using Switching Mechanism At 5’ End of RNA Template (SMARTer 5’) Rapid Amplification of cDNA Ends (RACE) technology (Takara Bio USA) adapted for antibodies to amplify the variable genes from heavy and kappa chains for each hybridoma. Briefly, RNA was extracted from hybridomas using QIAGEN RNeasy Mini Kit (QIAGEN, Valencia, CA), followed by first strand cDNA synthesis isotype-specific constant gene 3’ primers and the SMARTer II A Oligonucleotide and SMARTscribe reverse transcriptase. Amplification of cDNA was performed with SeqAmp DNA Polymerase (Takara), using a nested 3’ primer and a 5’ universal primer. Purified PCR products were then Sanger sequenced using 3’ constant gene primers (Azenta/GeneWiz, South Plainfield, NJ). Sequence results were blasted against the IMGT human databank of germline genes using V-Quest (http://imgt.org) and analyzed for clonality based on CDR3/junction identity and V(D)J usage. Unique clones from each clonal family were selected, and DNA was synthesized and cloned into in-house antibody expression vectors (pTWIST-hG1/hck) containing a human IgG1 constant region and kappa light chain constant region (Twist Biosciences, South San Francisco, CA).

### Generation of recombinant (r)mAbs in 293 suspension cells

DNA plasmids of the light and heavy chains were combined with Expifectamine1000 (Gibco) in a total of 24ml Opti-minimal essential media (MEM) and transfected in 3.5×10^6^cell/ml in a final volume of 200mlof Expi293 Expression Medium (Gibco). The next day, Enhancer 1 and Enhancer 2 were added (Gibco).

### ELISA

96-well plates (Immulon 4 HBX; Thermo Scientific) were coated with 50ul of H5 recombinant protein (2ug/ml) and incubated at 4°C overnight. The next day, 100ul of serially diluted IgG, starting at a concentration of 30ug/ml, was added to the plates in duplicate. Binding of the antibodies to the protein was evaluated using secondary antibody horseradish peroxidase (HRP)-labeled anti-human IgG (Sigma), diluted 1:3000. Positive control was anti-HA antibody^27^, while negative control was anti-SARS-CoV-2 spike antibody^13^.

### HI assay

First, a HA assay was performed using turkey RBCs to determine the viral HA titer. Next, 25ul of serially diluted IgG, starting at a concentration of 50ug/ml was incubated with with 8 HA units (HAU) of the virus for 1 hours (h) at room temperature (RT). Then, 50ul of 0.5% turkey RBCs was added, and the plates were placed at 4°C for 45min. Virus-negative wells were analyzed for pellet formation.

### *In vitro* microneutralization assay

Neutralization of clade 2.3.3.4b (H5N1-PR8) virus with human IgG was measured as previously described^28^. Serially diluted mAbs at a starting concentration of 30ug/ml were incubated in 100ul with 3.82×10^4^ TCID_50_ of the virus for 1h at RT followed by inoculation of MDCK cells plated on 96-well plates (Corning). After 48h of incubation at 37°C, an HA assay was performed to detect the presence of the virus. Positive control was anti-HA antibody^27^, while negative control was anti-SARS-CoV-2 spike antibody^13^.

### Neuraminidase inhibition (NI) assay

First, a neuraminidase assay of clade 2.3.3.4b (H5N1-PR8) virus was performed to determine the optimal virus concentration to be used in the NI assay. To measure the inhibitory activity of the antibodies, serially diluted mAbs (starting concentration 30ug/ml) were incubated in 50ul for 6h at 37 °C with an equal volume of the selected virus dilution in the fetuin-coated plates. The percentage of enzymatic inhibition was calculated relating the number in experimental wells to the mean number of virus only and wells with sample diluent only being negative (background) control. Half inhibitory concentration (IC_50_) was calculated based on dose-response curves, using top and bottom constraints of 0 and 100% GraphPad Prism 10.

### *In vivo* experiments

All experiments were performed according to protocols approved by the Icahn School of Medicine at Mount Sinai Institutional Animal Care and Use Committee (IACUC). Six-week-old female BALB/c mice were housed for at least one week before experiments (The Jackson Laboratory), and infections were performed on animals anesthetized with a ketamine-xylazine mixture. In prophylactic studies, mice were divided into four groups of five, receiving 5mg/kg, 1mg/kg, 0.3mg/kg, and 0.1mg/kg of anti-H5 mAbs intraperitoneally. An irrelevant antibody, an anti-SARS-CoV-2 spike mAb, was used to measure any non-specific protection. After 4h, mice were inoculated intranasally with 50% lethal doses (LD_50_) of A/bald eagle/FL/W22-134-OP/2022 (H5N1-A/PR/8/34 reassortant) virus in a total volume of 50ul PBS. Mice were monitored daily for body weight and mortality until day 14 post-infection or until all animals died, with those losing 25% or more of their initial body weight being euthanized. In therapeutic studies, three experimental groups of five mice were first infected with 5 LD_50_ of A/bald eagle/FL/W22-134-OP/2022 (H5N1-A/PR/8/34 reassortant) virus and mice were then treated at two, three and four days post-infection with 5mg/kg of anti-H5 mAbs and an irrelevant antibody, an anti-SARS-CoV-2 mAb served as control to measure any non-specific protection. Mice were monitored similarly daily for body weight and mortality. For viral clearance in lung analysis, as for prophylactic studies, groups of three mice received 5mg/kg of anti-H5 mAbs intraperitoneally, followed by intranasal inoculation with 5 LD_50_ of virus in 50ul PBS. Mice were euthanized on day 3 or 5 post-infection for lung titer analysis. Lungs were harvested and homogenized in 0.8ml of PBS using a BeadBlaster 24 (Benchmark) homogenizer. The homogenates were centrifuged (15min, 16,100 × *g*, 4°C) to remove cellular debris and stored in single-use aliquots at −80°C. Infectious virus titers were determined by plaque assay see supplementary material.

### Immune complexing and negative stain electron microscopy (EM)

Monoclonal antibody-binding fragment (Fab) and H5 protein (A/Jiangsu/NJ210/2023 or A/Red-tailed hawk/New York/NYCVH 22-8477/2022) were complexed at a 3:1 molar ratio for 1h at RT. For nsEM grids, 3ul of immune complexes were applied to glow-discharged, carbon-coated 400 mesh copper grids at a concentration of ∼10-20ug/ml (Electron Microscopy Services). Excess sample was blotted, and the complexes were stained with 2% w/v uranyl format for 60s twice. Imaging was performed on a Talos 200C or Tecnai Spirit T12 electron microscope. Micrographs were collected using Leginon. For each complex, 30k to 100k particles were picked and stacked using Appion and processed to generate 2D classes and 3D reconstructions using Relion 3.0 or Relion 4.0b1. Initial reference models were based on PDB: 6E7G and 4K62. Composite 3D reconstructions were made with UCSF ChimeraX.

### CryoEM grid preparation and imaging

Fabs of 12G1 and 20D10 were incubated with Jiangsu H5 and CR9114 Fab at a 3:1:3 molar ratio for 1h at RT prior to cryoEM grid preparation and placed on ice. Fabs of 1A1 and 6G1 were co-incubated with Jiangsu H5 and CR9114^27^ at a molar ratio of 3:3:1:3. A Vitrobot Mark IV (Thermo Fisher Scientific) was used to vitrify all samples at chamber temperature of 4°C, 100% humidity. To decrease orientation bias, immune complexes (∼0.7-0.85mg/ml) were mixed with n-Octyl-beta-D-glucoside (OBG, Anatrace) to a final OBG concentration of 0.1% before freezing. Datasets were collected on a 200 kV Glacios (Thermo Fisher) equipped with a Falcon IV direct electron detector. Data for the 12G1 complex and the 1A1 + 6G1 co-complex were collected using EPU at 190,000× magnification, with a pixel size of 0.725 Å and an exposure dose of 45 e−/Å². The 20D10 complex was collected similarly with a pixel size of 0.718 Å. Each dataset contained approximately 2,000 to 7,000 movie micrographs.

## Acknowledgements

Work in the Krammer laboratory was funded by institutional funding, NIAID Centers of Excellence for Influenza Research and Response (CEIRR, 75N93021C00014, reagent generation) and initial isolation of New York City H5N1 viruses was funded by Flu Lab. Work in the Ward laboratory was funded by NIAID Collaborative Influenza Vaccine Innovation Centers (CIVIC 75N39019C00051).

## Data Availability Statement

nsEM and cryoEM maps and models are deposited in the Electron Microscopy DataBank under accession IDs EMD-48510-48512, 48679-48690, 48692-48696, and in the Protein DataBank under accession IDs 9MQ1, 9MQ2, and 9MQ3.

## Author contributions statement

Study design: GPA, JAD and FK. Acquisition of data: GPA, ANL, TY, DB, ML, KB, PJMB, AJR, TJ, RW, CM, JH and JAD. Analysis and interpretation of data: GPA, ANL, TY. DB, ML, KB, PJMB, AJR, TJ, RW, CM, JH, ABW, JAD and FK. Drafting of the manuscript: GPA, JAD and FK. Study supervision: JAD and FK. Guarantor: FK. All authors reviewed the manuscript.

## Supplementary Material

### Cells

Madin-Darby canine kidney (MDCK) cells (ATCC #CCL-34) were passaged and cultured in Dulbecco’s modified Eagles medium (DMEM, Gibco) supplemented with 10% fetal bovine serum (FBS, Gibco) penicillin (100U/ml) and streptomycin (100ug/ml) (P/S) at 37°C and 5% CO_2_. MDCK cells were used for *in vitro* microneutralization assays. Expi293F cells were maintained in Expi293 Expression Medium in a shaking incubator at 37°C and 8% CO2. Expi293F cells were used for monoclonal antibody (mAb) production.

### Recombinant glycoproteins

HA proteins (as soluble trimers) and NA proteins (as soluble tetramers) were expressed from various virus strains, including H5N1 strains from birds and mammals, including humans. HA proteins used in this study are referred to with the use of GISAID numbers and GenBank numbers: HA was used as an antigen; A/mallard/New York/22-008760-007-original/2022 (H5) (GISAID number EPI2017135). HA protein of viruses isolated from New York Virus Hunters community science initiative (NYCVH)^29^ includes: A/chicken/New York/NYCVH 168127/2023 (H5N1) (GenBank number WPF52968.1), A/Canada goose/New York/NYCVH 23-453/2023 (H5N1) (GenBank number WPL83443.1), A/Canada goose/New York/NYCVH 22-9190/2022 (H5N1) (GenBank number WPF48542.1), A/peregrine falcon/New York/NYCVH 160820/2022 (H5N1) (GenBank number WPF49852.1), A/red-tailed hawk/New York/NYCVH 22-8477/2022 (H5N1) (GenBank number WPF47596.1). HA protein from the virus strain used to characterize the mAbs in vitro, A/bald eagle/FL/W22-134-OP/2022 (H5N1) (GISAID number EPI3661109). HA proteins from virus strains isolated from infected birds in 2014 include: A/chicken/Netherlands/14015531/2014 (H5N8) (GISAID number EPI2817295), A/northern pintail/WA/40964/2014 (H5N1) (GenBank number AJE30344.1). HA protein from a virus isolated from infected dairy cattle: A/dairy cattle/Texas/24-008749-001-original/2024 (GISAID number EPI3158678). HA proteins from virus strains isolated from infected humans: A/Vietnam/1203/2004 (H5N1) (GISAID number EPI1990181), A/Indonesia/5/2005 (H5N1) (GenBank number AFM78567.1), A/Shenzhen/1/2016 (H5N1) (GISAID number EPI687704), A/Cambodia/NPH230032_2/2023 (H5N1) (GISAID number EPI2419978), A/Wisconsin/67/2022 (H1N1) (GISAID number EPI2224788).

### Immunization

Eight to twelve-week-old H2L2 Harbour Mice^®^ immunized intraperitoneally with HA and NA from A/mallard/New York/22-008760-007-original/2022 (H5N1) adjuvanted with 40ug of poly I:C (InvivoGen, San Diego, CA). Each mouse received a prime followed by two boosts, and blood was collected two weeks after each boost to monitor the titer of serum antibodies via enzyme-linked immunosorbent assay (ELISA) and hemagglutination inhibition (HI) analysis. All experiments were performed according to protocols approved by the Icahn School of Medicine at Mount Sinai Institutional Animal Care and Use Committee (IACUC).

### Concentration and purification of recombinant (r)mAbs in 293 suspension cells

After seven days of transfecting Expi293F cells with the DNA plasmid of the light and heavy chains and incubation in a shaking incubator at 37°C, supernatant containing IgG was first clarified by centrifugation at 4,000g for 30 minutes (min), sing 0.45 μM Stericup filters (Sigma-Aldrich). The clarified supernatant was then purified using IgG-Fc packed columns (Thermo Scientific) by gravity flow, concentrated with Vivaspin 500 centrifugal concentrator, 30,000 molecular weight cut-off (MWCO) (GE Lifescience) and quantified with the nanodrop. The heavy and light chain plasmids of an irrelevant human IgG control monoclonal antibody were co-transfected and purified similarly.

### ELISA

96-well plates (Immulon 4 HBX; Thermo Scientific) were coated with 50ul of H5 recombinant protein (2ug/ml) and incubated at 4°C overnight. The next day, plates were blocked with 200ul/well of 3% non-fat milk in phosphate-buffered saline (PBS, Gibco) with 0.1% Tween (Fisher Scientific) (PBST) for 1hour (h) at room temperature (RT). After discarding the blocking solution, 100ul of serially diluted IgG, at a starting concentration of 30ug/ml in PBST-1% non-fat milk was added to the plates in duplicates for 2h at RT. Plates were washed three times with PBST, and mAbs binding was detected by a secondary horseradish peroxidase (HRP)-labeled anti-human IgG (GE Healthcare) diluted 1:3000 in PBST-1% non-fat milk, incubated for 1h at RT, followed by three PBST washes. 3,3′, 5,5′-tetramethylbenzidine (TMB, Thermo Scientific) was added for 10min, followed by Stop Solution (Invitrogen). Absorbance was determined at 450nm via a Synergy 4 (BioTek) plate reader. An anti-HA antibody, CR9114^27^ was used as a positive control, while an anti-severe acute respiratory syndrome coronavirus 2 (SARS-CoV-2) spike antibody^13^ was used as a negative control. Control wells contained only secondary antibody.

### Virus production

10-day-old embryonated chicken eggs (Charles River Laboratories) were inoculated with the virus and incubated at 37°C for 48h. Allantoic fluid was collected after the eggs were chilled at 4°C overnight and clarified by centrifugation at 3800 x g for 30min before freezing at -80°C.

### Determination of virus HA titer

HA titer of the viruses was determined in allantoic fluid collected from infected eggs. In brief, the collected allantoic fluid was spin at 3,000g for 10min, supernatant was collected and serially diluted (1:2) in a V-bottom 96-well plate (ThermoFisher), testing each dilution in replicates. Turkey red blood cells (RBCs) were first washed and then suspended in PBS, at a concentration of 0.5%, and were added to the virus-containing plate. After 45min of incubation at 4°C, virus positive wells were visualized based on the formation of a pellet. No pellet indicates binding of hemagglutinin to the sialic acid receptors on RBCs and hence, lack of a pellet was considered positive.

### *In vitro* microneutralization assay

To test neutralization of A/bald eagle/FL/W22-134-OP/2022 (H5N1-A/PR/8/34 reassortant), here referred to as clade 2.3.4.4b (H5N1-PR8) virus with human IgG, mAbs were serially diluted at a starting concentration of 30ug/ml in infection media (1xMEM with tosyl phenylalanyl chloromethyl ketone(TPCK)-treated trypsin), 3.82×10^4^ TCID_50_ of the virus was then added, and the mix was incubated in 100ul for 1h at RT, testing each antibody in duplicate. Virus-only controls were prepared by mixing virus with infection media. Virus-antibody mixes, virus-only mixes, or medium-only were then added to MDCK cells seeded at 25000cells/well in 96-well plates (Corning) the day prior. Following 1h incubation, cells were washed with PBS, and 100ul of infection medium was added to evaluate viral entry, while the mixed solutions of serially diluted antibodies in infection media were added to assess viral release. After 48h of incubation at 37°C, an HA assay was performed to detect the presence of the virus.

### Neuraminidase (NA) assay

The NA assay was performed for all virus stocks. Microtiter 96-well plates (Immulon 4 HBX; Thermo Fisher Scientific) were coated with 25ug/ml (100ul/well) of fetuin (Sigma) diluted in coating solution (KPL) and incubated overnight at 4°C. The following day, clade 2.3.4.4b (H5N1-PR8) virus was serially diluted 1:2 in PBS containing 5% bovine serum albumin (blocking solution) in a separate 96-well plate, starting with the undiluted stock. Fetuin-coated plates from the previous day were washed three times with PBS-T and viral dilutions in 100ul were transferred to the fetuin-coated plates and incubated at 37°C for 6h. The plates were washed four times with PBS-T. Peanut agglutinin conjugated to HRP (PNA-HRP; Sigma) at a concentration of 5ug/ml diluted in PBS (100ul/well) was added, and the reaction mixture was incubated at RT for 1.5h. After incubation, the plates were washed again four times with PBS-T and developed for 10min with 100ul/well TMB (Thermo Scientific). The developing process was stopped with 50ul/well 3M hydrochloric acid, and the reaction was immediately read at an optical density (OD) of 490nm using a Synergy H1 hybrid multimode microplate reader (BioTek). To determine the optimal virus concentration to be used in the neuraminidase inhibition (NAI) assay, the absorbance data were plotted in GraphPad Prism 7. The data were fitted to a nonlinear regression curve to determine the 50% effective concentration (EC_50_). The EC_50_ was subsequently used for NAI assays.

### Neuraminidase inhibition assay

Microtiter 96-well plates (Immulon 4 HBX; Thermo Fisher Scientific) were coated with 25ug/ml (100ul/well) of fetuin (Sigma) diluted in coating solution (KPL) and incubated overnight at 4°C. The following day, anti-H5 mAbs were serially diluted in sample dilution at a starting concentration of 30ug/ml in duplicate. As a positive control anti-NA antibody, 1G01 was used, and as a negative control anti-SARS-CoV-2 spike antibody. The clade 2.3.4.4b (H5N1-PR8) virus was diluted in PBS at 2x the determined EC_50_, in an NA-assay described previously. Fetuin-coated plates were washed with PBST, virus-antibody mixes, virus-only mixes, or medium only were added to the plates and incubated for 6h at 37°C. The plates were washed and incubated with 5µg/100µl/well of for 1.5h at RT. Plarted were washed four times with PBS-T and developed for 10min with TMB. The developing process was stopped with 3M hydrochloric acid, and the reaction was immediately read at an optical density (OD) of 490nm using a Synergy H1 hybrid multimode microplate reader (BioTek). For all mAbs, half inhibitory concentration (IC_50_) was calculated based on a dose-response curve, using top and bottom constraints of 0 and 100%, using GraphPad Prism 10.

### Infectious virus determination via plaque assay

Briefly, 250ul of 10-fold dilutions of lung homogenates in PBS were used to infect the day before plated MDCK cells and incubated for lh at 37°C, testing each dilution in duplicate. Cells were then washed once with PBS and overlayed with oxoid agar (Oxoid), serum-free 2×MEM/bovine serum albumin (BSA) containing diethylaminoethanol (DEAE) dextran, supplemented with (TPCK)-treated trypsin. After 48h of incubation, cells were fixed using 4% formaldehyde and stained for H5 antigens. Plaque forming units were determined by staining with a well-defined anti-H5 mAbs primary antibody, followed by goat anti-human IgG (Fab specific)-peroxidase antibody (Sigma-Aldrich) and positive cells were visualized with the use of TrueBlue substrate (KPL-Seracare).

### IgG digestion

Tris-ethylenediaminetetraacetic acid (EDTA), L-cysteine, and papain were combined at a final concentration of 100mM Tris, 2mM EDTA, 10mM L-cysteine, and 1mg/ml papain at 37°C for 15min to activate the enzyme. Purified IgG were incubated with activated papain at 37°C for 4-5h, using 40ul of activated papain per mg of IgG. Digestion was halted by adding iodoacetamide to a final concentration of 0.03M. 50-100ul of IgG Fc Capture Select (ThermoFisher) resin was washed with TBS and incubated with the digestion solution for 30min to 1h at RT to capture free Fc and undigested IgG. Resin was pelleted, and the supernatant was collected and concentrated using a 10kDa Amicon concentrator.

### Immune complexing and negative stain EM

Monoclonal Fab and H5 (either A/Jiangsu/NJ210/2023 or A/Red-tailed Hawk/New York/NYCVH 22-8477/2022) were complexed at a 3:1 molar ratio for 1h at RT. To prepare nsEM grids, 3u of immune complexes were applied to glow-discharged, carbon-coated 400 mesh copper grids at a concentration of ∼10-20ug/m (Electron Microscopy Services). The whatman filter paper was used to blot the excess sample, and 3ul of 2% w/v uranyl formate was applied to stain complexes for 60s twice. Samples were imaged on either a Talos 200C with a Falcon II direct electron detector and a CETA-16M camera (FEI) at 200 kV, 73,000 × magnification, and 2.00 Å/pixel; a Talos 200C with a Falcon II direct electron detector and a CETA 4k camera (FEI) at 200 kV, 73,000 × magnification, and 1.98 Å/pixel; or a Tecnai Spirit T12 (FEI) with a CMOS 4k camera (TVIPS) at 120 kV, 52,000 × magnification, and 2.06 Å/pixel. Micrographs were collected using Leginon. For each complex, 30k to 100k particles were picked and stacked using Appion and subsequently processed to generate 2D classes and 3D reconstructions using Relion 3.0 or Relion 4.0b1^30,31^. 3D reconstructions of A/Jiangsu/NJ210/2023 and A/Red-tailed Hawk/New York/NYCVH 22-8477/2022 complexes used initial reference models of 30 Å volumes generated from PDB 6E7G and 4K62, respectively. UCSF ChimeraX was used to generate composite 3D reconstructions^32^.

### CryoEM grid preparation and imaging

Fabs of 12G1 and 20D10 were incubated with Jiangsu H5 and CR9114^27^ Fab at a 3:1:3 molar ratio for 1h at RT prior to cryoEM grid preparation and placed on ice until ready for use. Fabs of 1A1 and 6G1 were co-incubated with Jiangsu H5 and CR9114 at a molar ratio of 3:3:1:3. Prior to sample application, a PELCO easiGlow (Ted Pella Inc.) was used to plasma treat Quantifoil R1.2/1.3 Cu, 300-mesh grids (Quantifoil Micro Tools GmbH) for 25s at 15mA. A Vitrobot Mark IV (Thermo Fisher Scientific) was used to vitrify all samples at chamber temperature of 4°C, 100% humidity, blot force 1, wait time 5s, and blot times between 3-5s. To decrease orientation bias, immune complexes (∼0.7-0.85 mg/mL) were mixed with n-Octyl-beta-D-glucoside (OBG, Anatrace) to a final OBG concentration of 0.1% immediately before freezing. Grids were screened and datasets were collected on a 200kV Glacios (Thermo Fisher) equipped with a Falcon IV direct electron detector. For the 12G1 complex and the 1A1 + 6G1 co-complex, automated data collection was executed using EPU (Thermo Fisher) at 190,000× nominal magnification and a pixel size of 0.725 Å, with an approximate exposure dose of 45 e-/Å2 442 and a nominal defocus range of -0.7 to -1.4um. The 20D10 complex was collected similarly, but with a pixel size of 0.718. Each dataset contained ∼2000-7000 movie micrographs.

### Cryo-EM data processing

The CryoSPARC Live software was used for image preprocessing, which automatically performs image motion corrections and initial CTF estimations^33^. Further data processing with processed micrographs was then performed in CryoSPARC. Blob picker was used for initial particle picking. After particle extraction particles were subjected to several iterations of 2D classification and 2D class selection to remove junk particles. A final stack of particles was then subjected to Ab-Initio Reconstruction. The ab-initio classes were then used as reference models for a Heterogeneous Refinement job. Particles corresponding to 3D volumes with the best resolution and alignment were then subjected to Homogeneous Refinement. To overcome preferred orientation issues, particles corresponding to orientations that were overrepresented were removed using Rebalance Orientations, and the remaining particles were used for a Non-uniform Refinement job.^34^ In the case of 20D10, another iteration of the steps from Heterogeneous Refinement to Non-uniform Refinement were performed to end up with a rebalanced and well-aligning particle subset. In the case of 6G1, the non-uniform refined particles were used for a Reference-Based Motion Correction job, after which the motion corrected particles were used for a second Non-uniform Refinement job. In all instances, non-uniform refined particles were symmetry expanded to the C3 axis in accordance with the trimeric nature of HA, and to maximize occupancy of the Fabs in the case of partial stoichiometry. The symmetry expanded particle set was then used for a 3D classification job using sphere masks around the epitope and Fab density. Highest-resolution and best-aligning classes were selected (in some cases aided by Regroup 3D Classes) and used for Local Refinement employing a soft solvent mask encompassing the HA trimer and the Fab. Following a Global CTF Refinement job, further Local Refinement was performed to obtain a high-resolution map. As the map of 20D10 still showed signs of orientation bias at this point, a Rebalance Orientations job followed by Local Refinement was performed to generate the final map.

### Atomic model building

The initial model of HA H5 Jiangsu was kindly shared by Olivia Swanson while initial models of 6G1, 12G1 and 20D10 Fab were generated using ABodyBuilder 2.0.^35^ Initial models of HA and Fabs were fitted into the map density using ChimeraX after which the complex was further refined manually using Coot^32,36^. Next several iterations of Real-space refinement in Phenix, followed by manual model building in Coot were performed to improve model statistics^37^. Model to map fit was validated using the integrated tools Molprobity and EMRinger in the Phenix software package.^38,39^ Orientations of all glycans were assessed and corrected using Privateer^40^. Due to diffuse density in the respective areas the following glycans/residues were not modeled: glycan N154 in the model of 20D10, glycan N154 in chain b in the model of 12G1, and residues 272 and 273 in chain A, B and C in the model of 6G1. Final atomic models were submitted to the Protein Data Bank with accession codes found in Table S1. Figures including atomic models were all produced using ChimeraX.

## Supplementary Figure

**Supplementary Figure 1.**
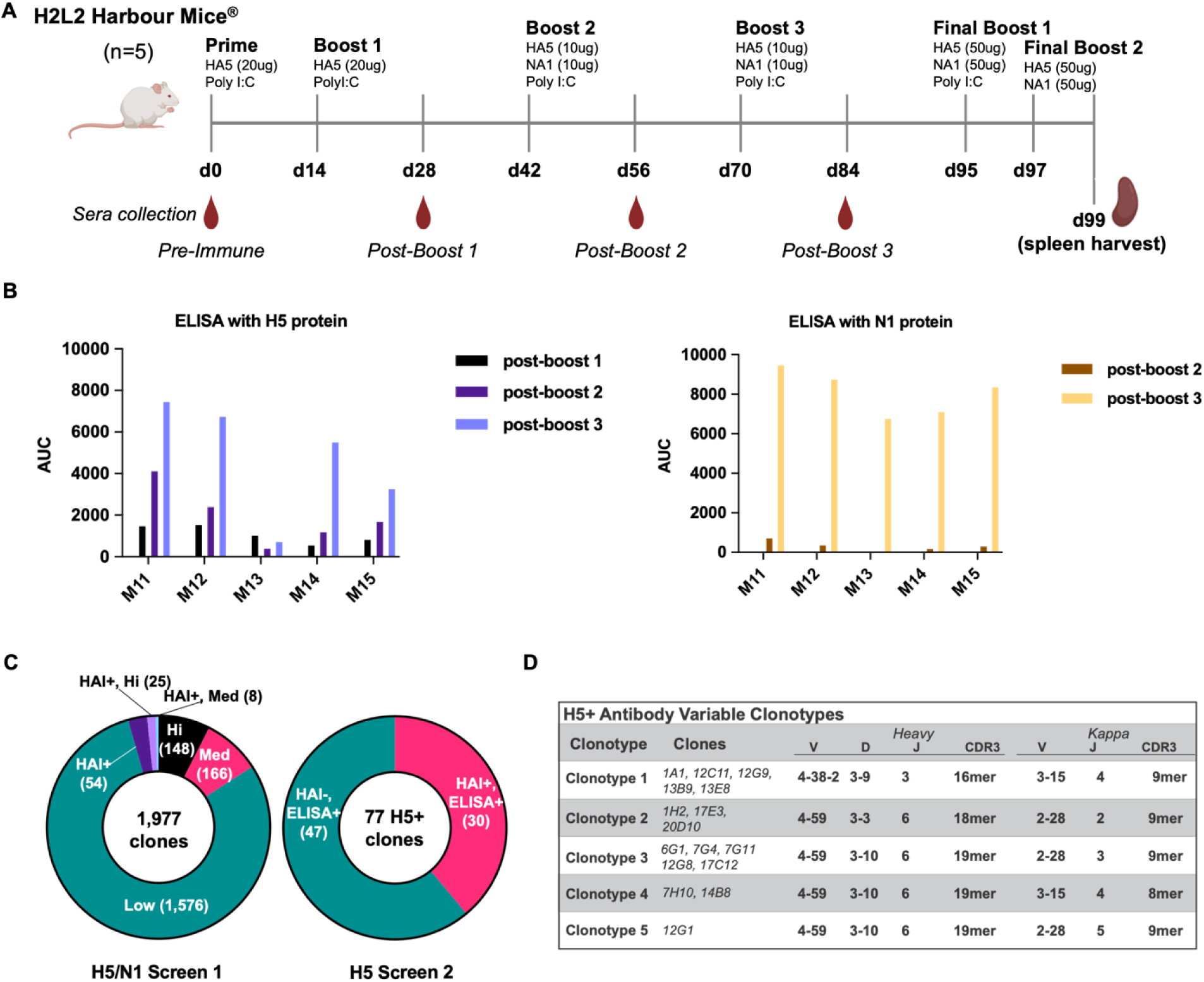
Immunization in H2L2 Harbour Mice^®^ with HA and NA protein of H5N1 clade 2.3.4.4b virus. **(A)** The timeline for vaccination is shown in the upper panel. Five female H2L2 Harbour Mice^®^ were immunized by intraperitoneal injection with the protein amounts shown in the figure, adjuvanted with poly IC. Sera was collected at the indicated time points, and the final immunization was performed two and four days before the spleen was harvested for hybridoma fusion. **(B)** Collected sera were tested against the H5 (left) and N1 (right) proteins by enzyme-linked immunosorbent assay (ELISA). Based on the results, a single mouse was selected and sacrificed for spleen harvest and hybridoma fusion. In this graph, the x-axis represents the individual mice, and the y-axis represents the area under the curve (AUC) for the response with the sera. The area under the curve (AUC) was calculated using GraphPad Prism 10, with the average plus three standard deviations of the blank wells serving as the baseline. **(C**) A total of 1,977 hybridoma clones were generated from the top responding mouse. Supernatants from the clones were first screened by ELISA coated with a combination of H5 and N1 proteins. Hemagglutinin inhibition (HI) assays were also performed at a fixed supernatant dilution of 1:10. ELISA binders were compared to a 1:100 dilution of unimmunized sera (normal mouse sera, NMS) and were classified as high (Hi) (>3x NMS), medium (Med) (1.5-3x NMS), and low (<1.5x NMS), with and without HI activity (right pie graph). All medium and high binding clones with and without HI activity were down-selected for secondary screening (401 clones) by H5-specific ELISAs and HI assays. Seventy-seven total clones (39%) rescreened positive by either ELISA or HI (left pie graph). The top thirty clones that were both ELISA and HI positive were down-selected for isotyping, variable segment sequencing and recombinant production as human G1/ kappa. **(D)** Clonotype analysis based on shared CDR3s (shared length and >90% amino acid identity) and shared VDJ/ VJ gene usage on the sequences of the top performing sixteen clones revealed five closely related clonotypes.

**Supplementary Figure 2.**
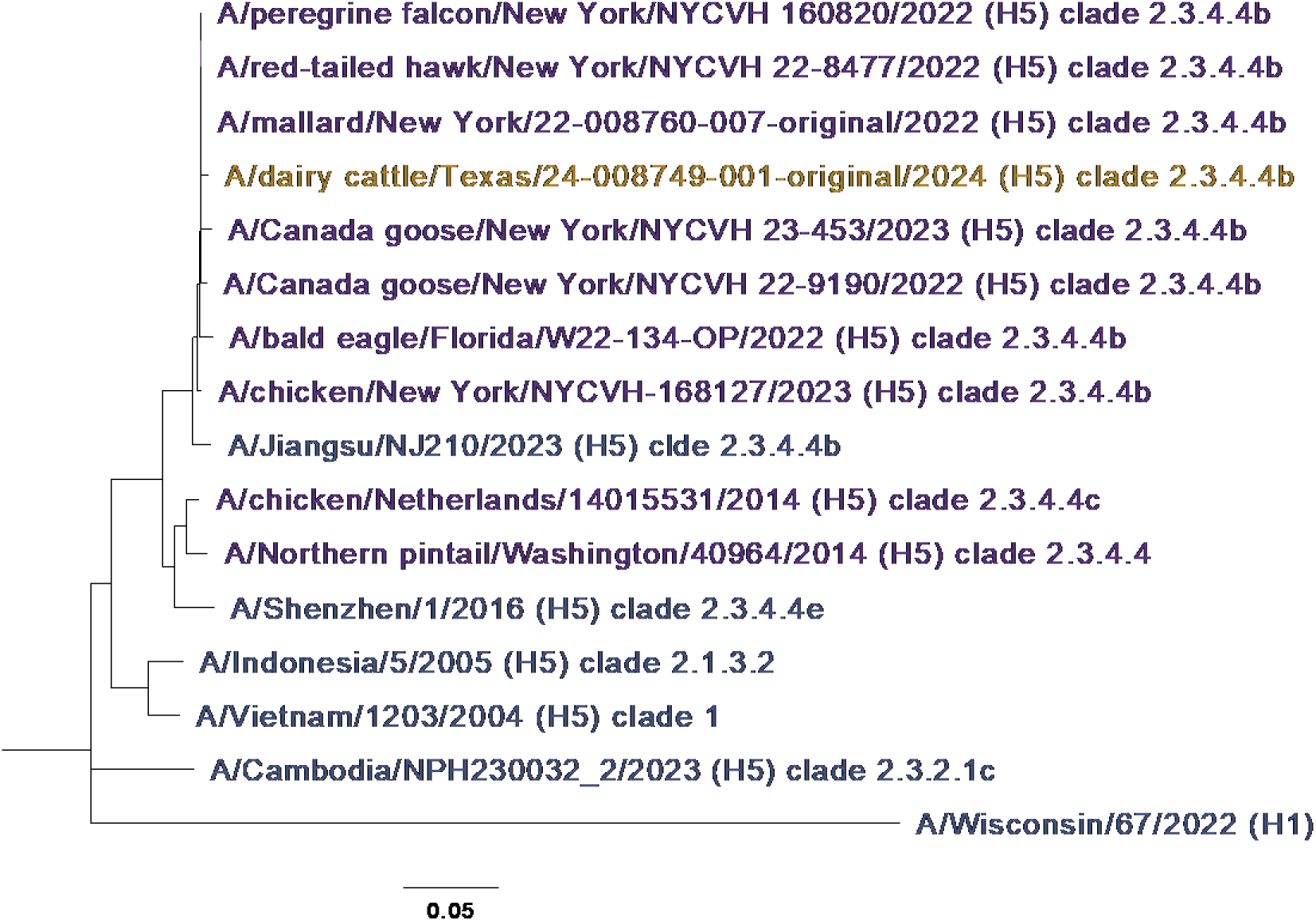
Phylogenetic analysis of the HA sequence of the viruses used throughout the study. The phylogenetic tree here was constructed using the HA amino acid (aa) sequence of H5N1 viruses isolated from infected birds and mammals, including humans. Color coded in purple are: the H5N1 virus used as the antigen, A/mallard/New York/22-008760-007-original/2022; the H5N1 virus strains obtained from the New York City Virus Hunters (NYCVH): A/chicken/New York/NYCVH 168127/2023, A/Canada goose/New York/NYCVH 23-453/2023, A/Canada goose/New York/NYCVH 22-9190/2022, A/peregrine falcon/New York/NYCVH 160820/2022, and A/red-tailed hawk/New York/NYCVH 22-8477/2022, the H5N1 virus strain used for cell culture assays, A/bald eagle/FL/W22-134-OP/2022; finally, H5N8 and H5N2 virus strains isolated in 2014 from two different geographic locations: A/chicken/Netherlands/14015531/2014 and A/Northern Pintail/Washington/40964/2014, respectively. Color coded in orange, is the recently detected H5N1 bovine virus: A/dairy cattle/Texas/24-008749-001-original/2024. Color coded in blue are virus strains isolated from humans infected with H5N1 virus strains from 2005 to 2023: A/Vietnam/1203/2004 (H5N1), A/Indonesia/5/2005 (H5N1), A/Shenzhen/1/2016 (H5N6), A/Cambodia/NPH230032_2/2023 (H5N1), A/Jiangsu/NJ210/2023 (H5N1) and virus strains isolated from humans infected with H1N1 virus A/Wisconsin/67/2022. The scale represents a 0.02% difference in amino acid sequence.

**Supplementary Figure 3.**
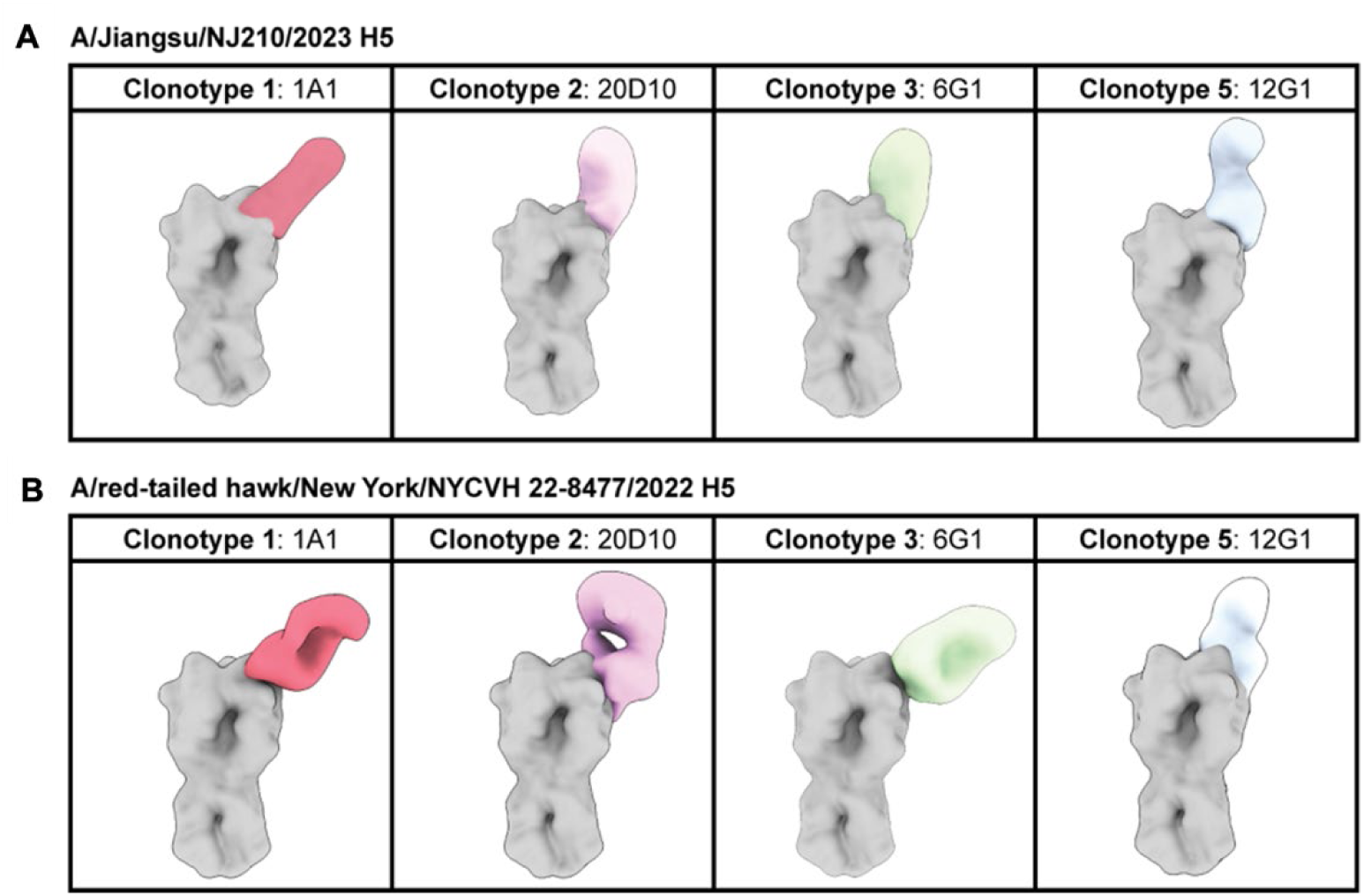
Negative stain EM reconstructions of representative mAbs for clonotypes 1-3 and 5. Maps demonstrate binding to either recombinant A/Jiangsu/NJ210/2023 H5 **(A)** or A/red-tailed hawk/New York/NYCVH 22-8477/2022 H5 **(B)** depicted on a map of A/Vietnam/1203/2004 H5 (PDB:6E7G) generated using ChimeraX.

**Supplementary Figure 4.**
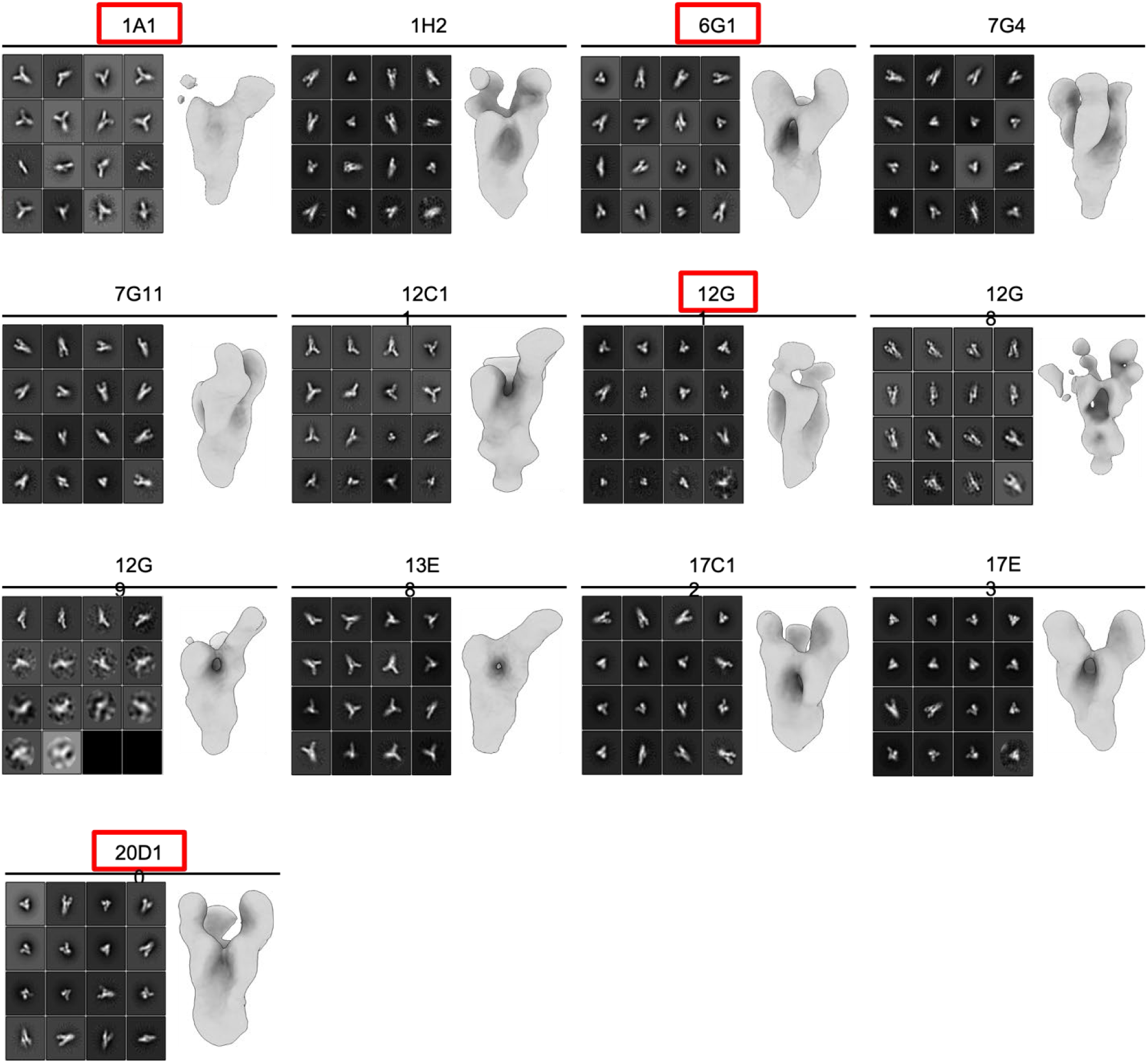
Negative stain EM representative 2D classes and 3D reconstructions of monoclonal antibodies bound to A/Jiangsu/NJ210/2023 H5. 2D class averages (left panels) representing particles used for negative stain 3D reconstructions (right panels) of mAbs in complex with recombinant A/Jiangsu/NJ210/2023 H5. Reconstructions were generated using Relion 3.0. mAbs selected as representative of their respective clonotypes and used for cryoEM analysis are boxed in red.

**Supplementary Figure 5.**
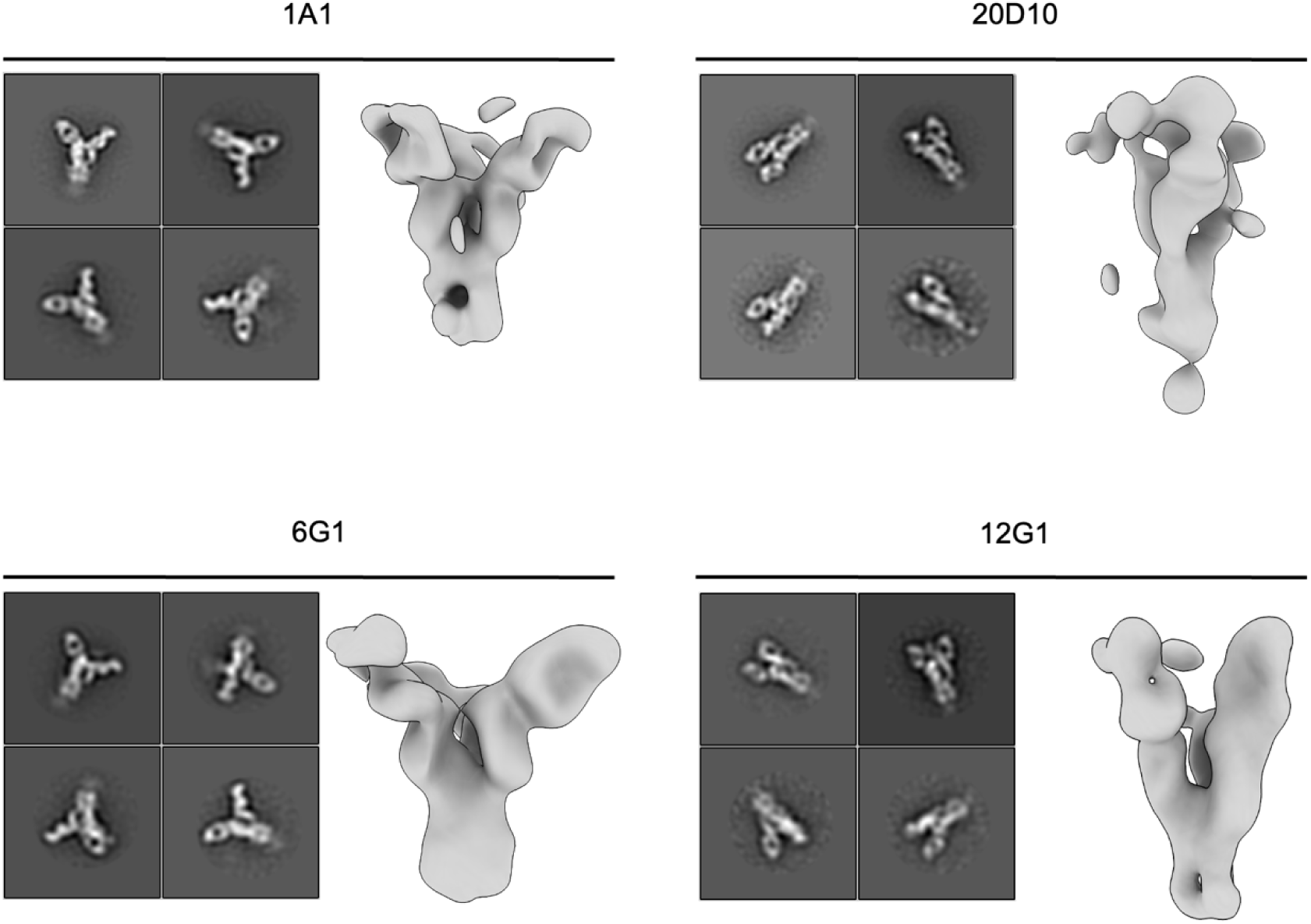
Negative stain EM representative 2D classes and 3D reconstructions of monoclonal antibodies bound to A/red-tailed hawk/New York/NYCVH 22-8477/2022 H5. 2D class averages (left panels) representing particles incorporated in negative stain 3D reconstructions (right panels) of mAbs in complex with recombinant A/Jiangsu/NJ210/2023 H5. Reconstructions were generated using Relion 3.0.

**Supplementary Figure 6.**
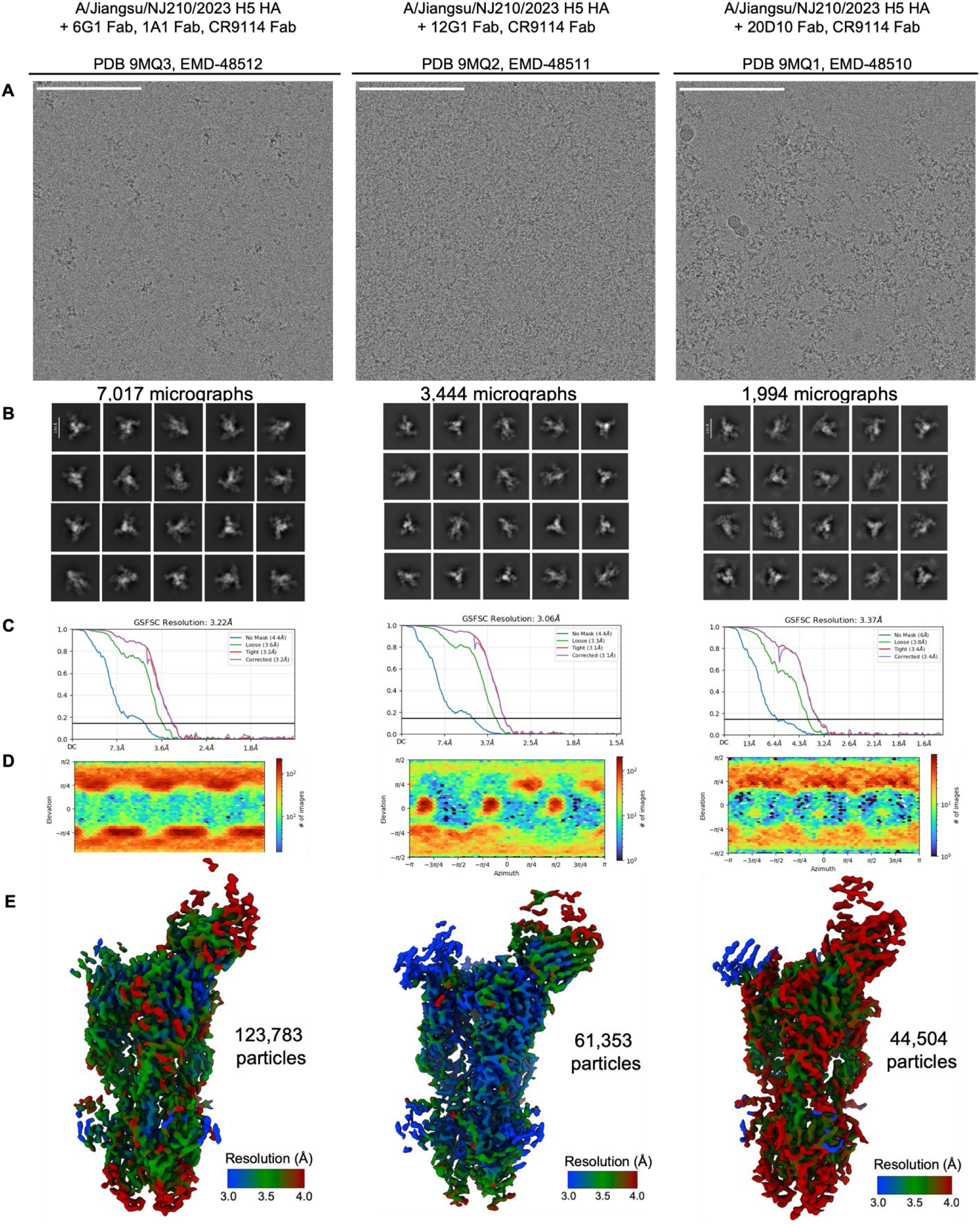
CryoEM workflow. **(A)** Representative micrographs with 100nm white scale bars. **(B)** Top 20 classes of final 2D class average selection after 2D clean up steps. **(C)** Fourier shell correlation (FSC) plots, **(D)** viewing distribution plots, **(E)** local resolution maps, and particle count for final 3D reconstructions.

**Supplementary Table 1.**
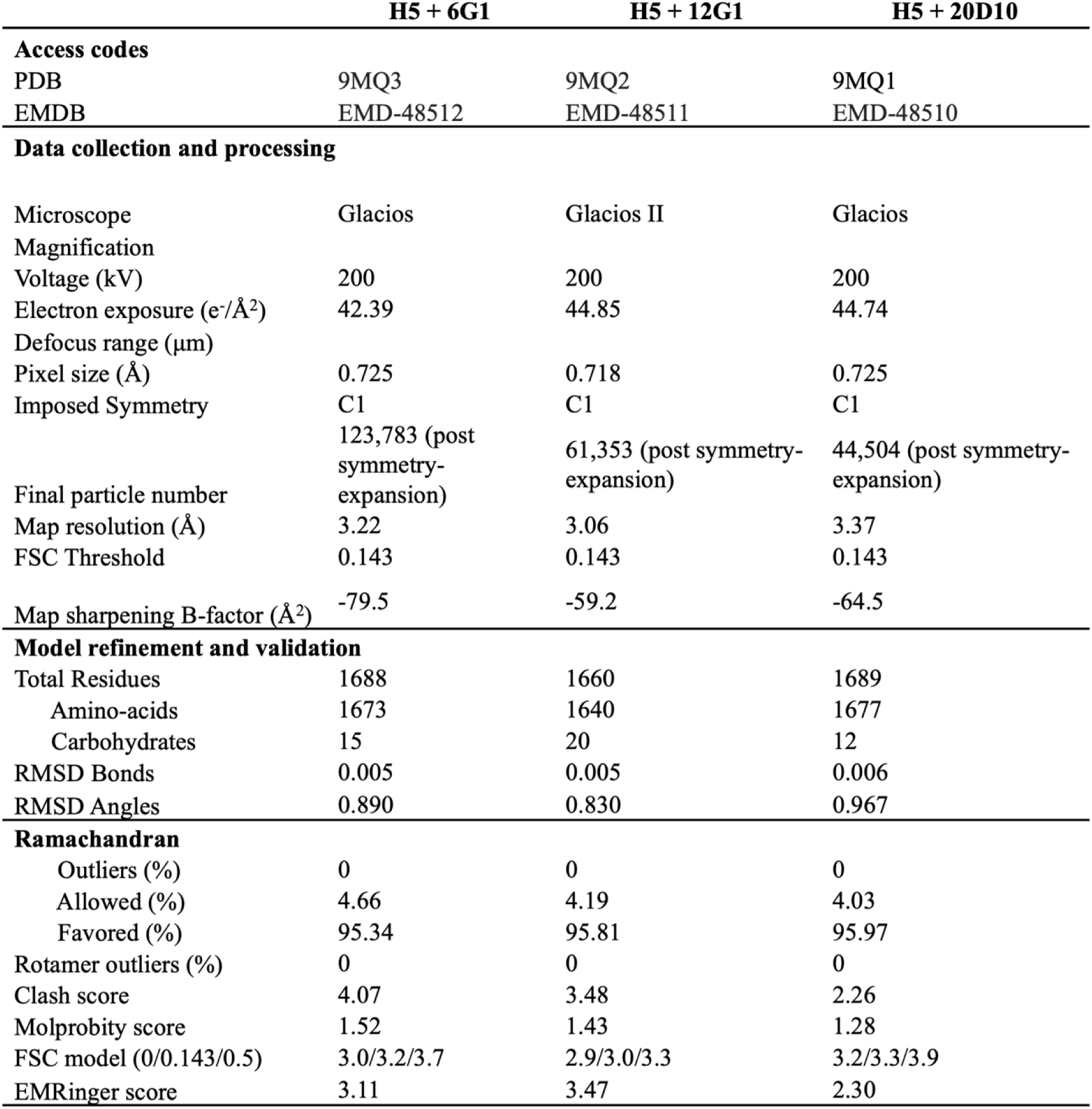
Cryo-EM data collection, processing, model refinement and validation statistics.

**Supplementary Table 2.**
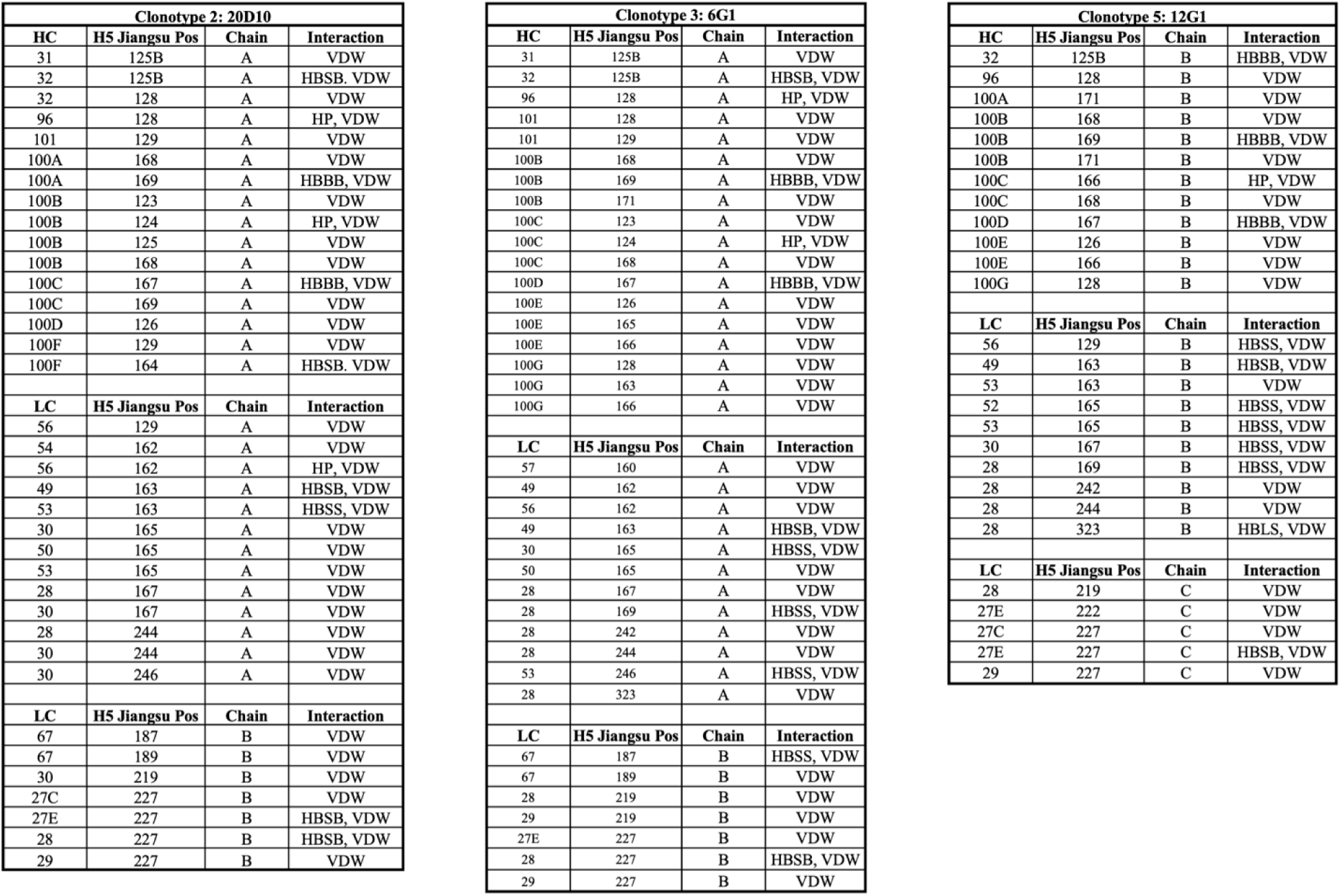
Contacts between monoclonal antibodies and A/Jiangsu/NJ210/2023 H5. The getcontacts program (https://getcontacts.github.io/index.html) was used on final models of 20D10, 6G1, and 12G1 in complex with A/Jiangsu/NJ210/2023 H5 to predict protein-protein interactions between amino acid residues. Relevant interactions are abbreviated as follows: van der Waals (VDW), side-chain to backbone hydrogen bonding (HBSB), backbone to backbone hydrogen bonding (HBBB), side-chain to side-chain hydrogen bonding (HBSS), and hydrophobic (HP).

## References

1 Sims, L. D. et al. Avian influenza in Hong Kong 1997-2002. Avian Dis 47, 832–838 (2003). 10.1637/0005-2086-47.s3.832

2 Webby, R. J. & Uyeki, T. M. An Update on Highly Pathogenic Avian Influenza A(H5N1) Virus, Clade 2.3.4.4b. J Infect Dis 230, 533–542 (2024). 10.1093/infdis/jiae379

3 US Department of Agriculture Animal and Plant Health Inspection Service. Confirmations of highly pathogenic avian influenza in commercial and backyard flocks. 2024. https://www.aphis.usda.gov/livestock-poultry-disease/avian/avian-influenza/hpai-detections/commercial-backyard-flocks. Accessed 20 June 2024

4 Plaza, P. I., Gamarra-Toledo, V., Eugui, J. R. & Lambertucci, S. A. Recent Changes in Patterns of Mammal Infection with Highly Pathogenic Avian Influenza A(H5N1) Virus Worldwide. Emerg Infect Dis 30, 444–452 (2024). 10.3201/eid3003.231098

5 US Department of Agriculture Animal and Plant Health Inspection Service. Federal and State Veterinary Agencies Share Update on HPAI Detections in Oregon Backyard Farm, Including First H5N1 Detections in Swine. 2024. https://www.aphis.usda.gov/news/agency-announcements/federal-state-veterinary-agencies-share-update-hpai-detections-oregon. Accessed 4 December 2024.

6 Centers for Disease Control and Prevention. A(H5N1) Bird Flu Response Update November 18, 2024. 2024. https://www.cdc.gov/bird-flu/spotlights/h5n1-response-11152024.html. Accessed 18 November 2024.

7 Adisasmito, W. et al. Effectiveness of antiviral treatment in human influenza A(H5N1) infections: analysis of a Global Patient Registry. J Infect Dis 202, 1154–1160 (2010). 10.1086/656316

8 Wang, C. et al. A human monoclonal antibody blocking SARS-CoV-2 infection. Nat Commun 11, 2251 (2020). 10.1038/s41467-020-16256-y

9 Focosi, D. et al. Monoclonal antibody therapies against SARS-CoV-2. Lancet Infect Dis 22, e311–e326 (2022). 10.1016/S1473-3099(22)00311-5

10 Widjaja, I. et al. Towards a solution to MERS: protective human monoclonal antibodies targeting different domains and functions of the MERS-coronavirus spike glycoprotein. Emerg Microbes Infect 8, 516–530 (2019). 10.1080/22221751.2019.1597644

11 Meade, P. S. et al. Detection of clade 2.3.4.4b highly pathogenic H5N1 influenza virus in New York City. J Virol 98, e0062624 (2024). 10.1128/jvi.00626-24

12 Stadlbauer, D. et al. Antibodies targeting the neuraminidase active site inhibit influenza H3N2 viruses with an S245N glycosylation site. Nat Commun 13, 7864 (2022). 10.1038/s41467-022-35586-7

13 Clark, J. J. et al. Protective effect and molecular mechanisms of human non-neutralizing cross-reactive spike antibodies elicited by SARS-CoV-2 mRNA vaccination. Cell Rep 43, 114922 (2024). 10.1016/j.celrep.2024.114922

14 World Health Organization. Assessment of risk associated with recent influenza A(H5N1) clade 2.3.4.4b viruses. 2022. https://cdn.who.int/media/docs/default-source/influenza/avian-and-other-zoonotic-influenza/h5-risk-assessment-dec-2022.pdf. Acessed 21 December 2022.

15 Centers for Disease Control and Prevention. CDC Confirms First Severe Case of H5N1 Bird Flu in the United States. 2024. https://www.cdc.gov/media/releases/2024/m1218-h5n1-flu.html. Accessed 18 December 2024.

16 Jassem, A. N. et al. Critical Illness in an Adolescent with Influenza A(H5N1) Virus Infection. N Engl J Med (2024). 10.1056/NEJMc2415890

17 Castillo, A. et al. The first case of human infection with H5N1 avian Influenza A virus in Chile. J Travel Med 30 (2023). 10.1093/jtm/taad083

18 Bruno, A. et al. First case of human infection with highly pathogenic H5 avian Influenza A virus in South America: A new zoonotic pandemic threat for 2023? J Travel Med 30 (2023). 10.1093/jtm/taad032

19 Kilbourne, E. D. Influenza pandemics of the 20th century. Emerg Infect Dis 12, 9–14 (2006). 10.3201/eid1201.051254

20 Andreev, K. et al. Genotypic and phenotypic susceptibility of emerging avian influenza A viruses to neuraminidase and cap-dependent endonuclease inhibitors. Antiviral Res 229, 105959 (2024). 10.1016/j.antiviral.2024.105959

21 Gu, C. et al. A human isolate of bovine H5N1 is transmissible and lethal in animal models. Nature 636, 711–718 (2024). 10.1038/s41586-024-08254-7

22 Tan, S. K. et al. A Randomized, Placebo-Controlled Trial to Evaluate the Safety and Efficacy of VIR-2482 in Healthy Adults for Prevention of Influenza A Illness (PENINSULA). Clin Infect Dis 79, 1054–1061 (2024). 10.1093/cid/ciae368

23 Stadlbauer, D. et al. Broadly protective human antibodies that target the active site of influenza virus neuraminidase. Science 366, 499–504 (2019). 10.1126/science.aay0678

24 Margine, I., Palese, P. & Krammer, F. Expression of functional recombinant hemagglutinin and neuraminidase proteins from the novel H7N9 influenza virus using the baculovirus expression system. J Vis Exp, e51112 (2013). 10.3791/51112

25 Amanat, F., Meade, P., Strohmeier, S. & Krammer, F. Cross-reactive antibodies binding to H4 hemagglutinin protect against a lethal H4N6 influenza virus challenge in the mouse model. Emerg Microbes Infect 8, 155–168 (2019). 10.1080/22221751.2018.1564369

26 Duty, J. A. et al. Discovery and intranasal administration of a SARS-CoV-2 broadly acting neutralizing antibody with activity against multiple Omicron subvariants. Med 3, 705–721 e711 (2022). 10.1016/j.medj.2022.08.002

27 Dreyfus, C. et al. Highly conserved protective epitopes on influenza B viruses. Science 337, 1343–1348 (2012). 10.1126/science.1222908

28 Nachbagauer, R. et al. Defining the antibody cross-reactome directed against the influenza virus surface glycoproteins. Nat Immunol 18, 464–473 (2017). 10.1038/ni.3684

29 Marizzi, C. & Wright, L. Hunting emerging viruses through participatory community science. Nat Microbiol 9, 578–581 (2024). 10.1038/s41564-024-01604-1

30 Lander, G. C. et al. Appion: an integrated, database-driven pipeline to facilitate EM image processing. J Struct Biol 166, 95–102 (2009). 10.1016/j.jsb.2009.01.002

31 Kimanius, D., Dong, L., Sharov, G., Nakane, T. & Scheres, S. H. W. New tools for automated cryo-EM single-particle analysis in RELION-4.0. Biochem J 478, 4169–4185 (2021). 10.1042/BCJ20210708

32 Pettersen, E. F. et al. UCSF ChimeraX: Structure visualization for researchers, educators, and developers. Protein Sci 30, 70–82 (2021). 10.1002/pro.3943

33 Punjani, A., Rubinstein, J. L., Fleet, D. J. & Brubaker, M. A. cryoSPARC: algorithms for rapid unsupervised cryo-EM structure determination. Nat Methods 14, 290–296 (2017). 10.1038/nmeth.4169

34 Punjani, A., Zhang, H. & Fleet, D. J. Non-uniform refinement: adaptive regularization improves single-particle cryo-EM reconstruction. Nat Methods 17, 1214–1221 (2020). 10.1038/s41592-020-00990-8

35 Abanades, B. et al. ImmuneBuilder: Deep-Learning models for predicting the structures of immune proteins. Commun Biol 6, 575 (2023). 10.1038/s42003-023-04927-7

36 Emsley, P. & Cowtan, K. Coot: model-building tools for molecular graphics. Acta Crystallogr D Biol Crystallogr 60, 2126–2132 (2004). 10.1107/S0907444904019158

37 Liebschner, D. et al. Macromolecular structure determination using X-rays, neutrons and electrons: recent developments in Phenix. Acta Crystallogr D Struct Biol 75, 861–877 (2019). 10.1107/S2059798319011471

38 Barad, B. A. et al. EMRinger: side chain-directed model and map validation for 3D cryo-electron microscopy. Nat Methods 12, 943–946 (2015). 10.1038/nmeth.3541

39 Chen, V. B. et al. MolProbity: all-atom structure validation for macromolecular crystallography. Acta Crystallogr D Biol Crystallogr 66, 12–21 (2010). 10.1107/S0907444909042073

40 Agirre, J. et al. Privateer: software for the conformational validation of carbohydrate structures. Nat Struct Mol Biol 22, 833–834 (2015). 10.1038/nsmb.3115

